# Polystyrene Nanoplastics Accumulate in Murine Cortex and Induce Transient Microglial Activation via Endolysosomal Retention

**DOI:** 10.64898/2026.03.24.712727

**Authors:** Alireza Tavakolpournegari, Unikrishnan Kannan, Mary Gregory, Julie Dufresne, Santiago Costantino, Stephane Lefrancois, Daniel G. Cyr

## Abstract

Environmental degradation and accumulation of plastics results in micro- and nanoplastics (MNPLs) that are small enough to cross biological barriers, including the blood–brain barrier. Microglia, resident immune cells of brain, are critical regulators of neuroimmune homeostasis and represent a cellular target of nanoplastic exposure. In this study, we assessed the neurotoxic effects of two sizes of polystyrene nanoplastics (PS-NPs; 100 nm and 500 nm) using integrated *in vivo* and *in vitro* exposure and washout paradigms. *In vivo* exposure in mice (60 days; 0.15 or 1.5 mg/day) showed the accumulation of both PS-NP sizes in the cerebral cortex without histopathological damage. However, cortical microglia showed pronounced morphological remodeling, observed as increased expression of Iba1 and GFAP. Transcriptomic profiling of cortical tissue revealed a strong size-dependent response. The 100 nm PS-NP group revealed 18 DEGs (|log_2_FC| ≥ 2, padj < 0.05), whereas the 500 nm PS-NPs showed more than 4,000 DEGs, including upregulation of immune- and microglia-associated genes (CCL5, CXCL10, LCN2, LYZ2) and downregulation of synaptic and neuronal signaling genes (GRIN2B, SYN1, STX1B, MAP1B, ITPR1/2). *In vitro* assessment, using BV2 microglia cells, showed internalization of PS-NPs via the endolysosomal pathway, with strong co-localization to Rab7- and LAMP2-positive compartments and prolonged intracellular retention following exposure washout. Also, microglial activation markers (Iba1, CD68) exhibited a transient, size- and concentration-dependent increase, correlated with intracellular particle burden rather than cumulative exposure. Overall, these findings demonstrate that PS-NPs accumulate in brain, driving size-dependent microglia activation and transcriptomic reprogramming, even after cessation of exposure to PS-NPs.

**Highlights:**

- PS-NPs (100 nm and 500 nm) reach mouse cerebral cortex following 60-day oral exposure.
- PS-NPs were internalized by microglia; accumulated in endolysosomal compartments.
- PS-NP exposure induced transient microglial activation without sustained cytotoxicity.
- Microglial activation was correlated with intracellular PS-NPs burden.
- Transcriptomics revealed disruption of neuroimmune and microglial regulatory pathways.

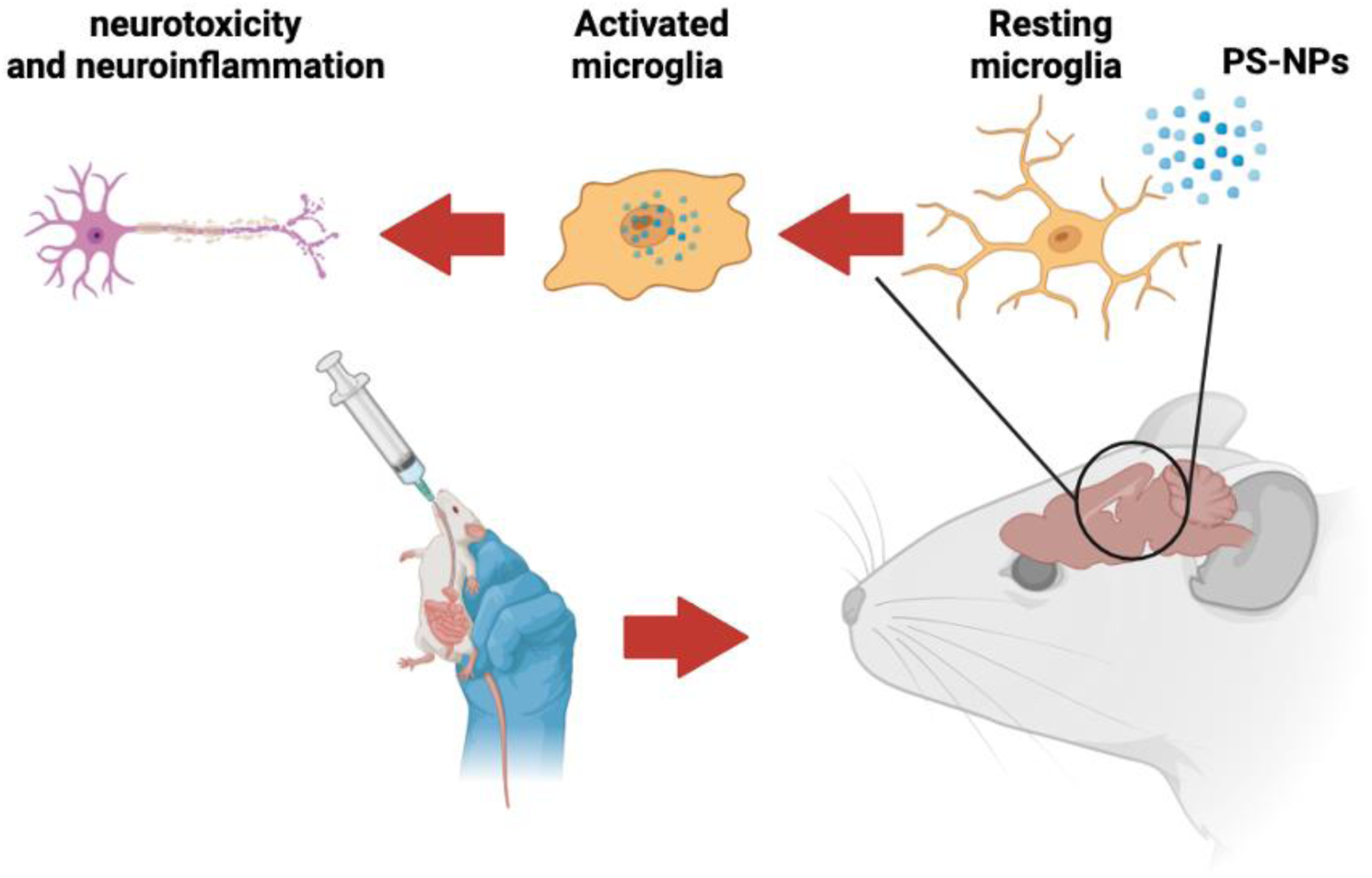

## 1. Introduction

Plastic pollution is one of the most pressing global environmental issues, driven by the accelerating growth of plastic manufacturing and the persistent accumulation of plastic waste in natural environments [1]. In 2023, worldwide plastic production was estimated to reach approximately 436 million metric tons, bringing the total metric tons of plastic manufactured to date at over 8 billion. Only about 9% of this production is recycled, with the remainder being either incinerated, buried in landfills, or released into the environment [2]. Once released, plastics undergo degradation and fragmentation into micro- and nanoplastics (MNPLs), which are small enough to enter and accumulate within the cells of target organs [3]. Recent estimates suggest that 0.1-5 grams of plastic per week might be ingested by humans through food, water, and air contamination [4]. Due to their remarkable bioavailability, increasing scientific attention has been directed towards MNPLs, their presence in human tissues, and their capacity to cross critical biological barriers, including the blood–brain barrier (BBB) [5].

Studies have shown that the brain is a target for the accumulation of MNPLs [6]. Emerging scientific evidence indicates that MNPLs may induce neuroinflammation, disturb neuronal function, and enhance the risk of neurodegenerative diseases [7]. Recent observations, including the highly publicized finding of what was described as “a plastic brain” underscore the growing concern that the MNPLs of various sizes and polymer types can accumulate in the central nervous system [8]. The role of glial cells in MNPLs neurotoxicity has gained particular attention due to their multifunctional role in brain homeostasis [9, 10]. Microglia are resident immune cells of the central nervous system and play crucial roles in maintaining brain homeostasis, regulating neuronal function, and orchestrating neuroimmune responses [11]. Alterations in microglial physiology may have profound implications for central nervous system health.

Studies have reported that MNPLs directly interact with microglia. These result in morphological changes, activation, release of pro-inflammatory cytokines (IL-1β, IL-1), induction of oxidative stress, mitochondrial dysfunction, and impairment of the autophagy–lysosomal pathways [12–14]. *In vivo* models of MNPLs exposure similarly exhibit cognitive deficits, behavioral abnormalities, and memory impairments consistent with microglia-mediated neurotoxicity [7, 9]. Such findings highlight microglia as cellular targets of MNPLs and emphasize the need to understand how plastic-derived particles modulate neuroimmune regulation and contribute to increased risk of neurodegenerative diseases.

Despite these advances, most existing studies focus primarily on exposure-outcome relationships and provide limited insights into the dynamic mechanisms by which MNPLs of different sizes and polymer types interact with microglia. In particular, little is known regarding the response of microglia to polystyrene nanoplastics (PS-NPs) under different exposure conditions, *in vivo* and *in vitro*, and whether or not PS-NPs-induced alterations persist after cessation of exposure to PS-NPs. Such observations in other cell types have highlighted the fact that PS-NPs-induced toxicity is not limited to acute interactions but may involve chronic, dose-cumulative effects, reinforcing the need to investigate post-exposure persistence and long-term cellular responses [15]. Addressing these gaps in knowledge is essential for understanding the real-world relevance of MNPL neurotoxicity.

In this study we investigated the neurotoxicity of two sizes of PS-NPs (100 and 500 nm), representative of environmental nanoplastics commonly found in food packaging using both *in vivo* and *in vitro* approaches.

## 2. Materials and Methods

### 2.1. Characterization of Polystyrene nanoplastics (PS-NPs)

Non-fluorescent and fluorescent PS-NPs (100 and 500 nm) were purchased from Alpha Nanotech Inc. (Vancouver BC). The characterization of PS-NPs (hydrodynamic and dry state size, Z-potential etc.) were supplied by the manufacturer.

### 2.2. Animal handling and PS-NPs exposure

Adult C57BL/6 male mice (48–50 days of age) were obtained from Charles River Laboratories (St Constant, QC) and acclimated for 5–7 days prior to experimentation.

Animals were housed under a 12-h light/12-h dark cycle with *ad libitum* access to food and water. Mice received a daily oral gavage (0.2 mL) of PS-NPs (100 or 500 nm) in sterile H_2_O for 60 consecutive days at a dose of 0.15 or 1.5 mg/day. The 60-day regimen corresponds to 1.5 cycles of spermatogenesis, a design frequently incorporated into reproductive toxicology studies. The dose was selected based on recent studies using comparable regimens and exposure conditions and based on estimates of human-equivalent intake after interspecies scaling [4]. Oral gavage was chosen as ingestion is considered the major route of PS-NP [12, 16]. Vehicle control mice were gavaged with sterile H_2_O alone. Body weights were recorded once weekly and animals were monitored daily for general health and behavior. On day 60, mice were anesthetized with isoflurane and euthanized by cervical dislocation under deep anesthesia. For experiments using fluorescently labeled PS-NPs, mice were administered PS-NPs daily via gavage, as described previously. Mice were sampled at 24 h (day 1) and 7 days (day 8) following the first PS-NPs administration. Euthanasia and tissue collection were conducted in accordance with protocols approved by the University Animal Care Committee.

### 2.3. Histology

Following euthanasia, the brain was immediately removed from each animal, rinsed briefly in sterile PBS, blotted dry, and fixed in 10% neutral buffered formalin. After fixation, samples were processed through graded alcohols, xylene and embedded in paraffin for histological analysis. Sections were prepared by the Histology Core (Goodman Cancer Institute, McGill University, Montreal, QC) and visualization was done using a Zeiss LSM790 confocal microscope (Carl Zeiss Microscopy GmbH, Germany). For histological evaluation, paraffin-embedded murine cerebral cortex sections were first deparaffinized by immersion in xylene, then rehydrated in graded ethanol (100%, 95%, 90%, 80%, 70%, and 0%) for 5 min each. The sections were stained with hematoxylin to visualize nuclei, followed by eosin counterstaining to delineate cytoplasmic and extracellular tissue architecture. After staining, sections were dehydrated through an ascending graded ethanol series, cleared in xylene, and mounted in Permount (Fisher Scientific, Ottawa, ON). Prepared slides were subsequently examined and imaged under a bright-field light microscope.

#### 2.4.1. Immunofluorescence

To investigate the effects of PS-NPs on microglia and astrocytes, two specific markers were used: Iba1 (Ionized calcium-binding adaptor molecule 1) and GFAP (Glial fibrillary acidic protein). Iba1 is a cytoplasmic protein specifically expressed in microglia and plays a critical role in actin cytoskeleton remodeling, phagocytosis, and inflammatory activation [17]. GFAP is the principal intermediate filament of astrocytes and is an essential structural and functional constituent of cortical glial cells [18].

Sections (5 µm) were deparaffinized by immersion in xylene (3 × 5 min) and rehydrated through a series of graded of ethanol to a final neutral solution of glycine (0.3 M). Heat-induced epitope retrieval was performed in citrate buffer (pH 6.0) for 10 min, followed by cooling to room temperature for ≥30 min and rinsing in distilled water. Sections were then equilibrated in TBST with 0.3 M glycine, blocked in TBST containing BSA (5%) and goat serum (5%) for 30 min at 37 °C, and incubated with primary antibodies against Iba1 (rabbit polyclonal, 2 μg/mL; #019-19741, Fujifilm, Wako, TX) or GFAP (rabbit monoclonal, 0.104 μg/mL; #ab68428, Abcam, Waltham, MA) overnight at 4 °C. Sections were then washed in TBST (3 × 5 min), and incubated with Alexa Fluor® 488-conjugated anti-rabbit secondary antibody (1 µg/mL; #111-545-144, Jackson Immuno Research, West Grove, PA) for 2 h at room temperature. Nuclei were stained with Hoechst dye (Invitrogen, Burlington ON). Slides were washed in TBST with glycine, mounted with Fluoromount (Southern BioTech, Birmingham, AL) and imaged as Z-stacks on a Zeiss LSM780 confocal microscope. Microglial and astrocyte morphology were quantified using Fiji/ImageJ [19] with identical processing parameters across different groups using skeleton and Sholl analyses.

#### 2.4.2. Skeleton analysis

Skeleton analysis was conducted to quantify specific parameters of microglial morphology, including branch length, number of branches, and junctional points. A median filter was then applied to remove background noise. Consistent thresholding was achieved using the Otsu method (*Image > Adjust > Threshold*) across all samples [19].

The thresholded images were converted into binary form and skeletonized (*Process > Binary > Skeletonize*) to reduce microglial structures to one-pixel-wide representations. Quantitative measurements were then obtained using the *Analyze Skeleton* plugin, which computed parameters such as total branch length, shortest and longest branch segments, and the number of branching junctions.

#### 2.4.3. Sholl analysis

Sholl analysis was performed to evaluate the extent of microglial branching and overall morphological complexity. Each image was initially selected and duplicated to preserve the integrity of the original data. To minimize background interference, a median filter was applied through the *Process* function. Subsequently, a consistent threshold was applied across all images using the Otsu method (*Image > Adjust > Threshold*), ensuring uniform image segmentation.

Following thresholding, the region of interest encompassing individual microglial cells was selected. The *Sholl Analysis* option within the Simple Neurite Tracer plugin was employed to quantify the branching intersections. The analysis generated both a Sholl image and a corresponding Sholl profile, which represented the frequency of intersections as a function of radial distance from the soma. This approach provided a quantitative measurement of dendritic arbor complexity and microglial process distribution.

### 2.5. RNA Extraction and Purification

Mouse brains from all groups (control, 100 nm and 500 nm PS-NPs) were collected at the time of euthanasia and placed on a sterile weighing dish on ice. The cortex of each brain was carefully excised, frozen immediately in liquid nitrogen, and subsequently stored at -80°C until further processing. RNA was extracted using a Macherey-Nagel Nucleospin RNAPlus kit (D-Mark, Mississauga ON) according to manufacturer’s instructions, with the following modifications: The volume of lysis buffer (500 μl/350 mg of tissue) was added to a 1.5 ml microcentrifuge tube. The tissue was then ground within the sample tube using a sterile mini pestle (Fisher Scientific). The tube was vortexed briefly and then returned to dry ice and to the -80°C freezer until further processing. The samples were then centrifuged at 13 000 g for 1 min. Prior to RNA elution, 0.5 µl of RiboLock (Thermo Fisher Scientific, Waltham, MA, USA) was added to each tube. RNA was then eluted with RNase-free water and immediately placed on ice. RNA concentrations were measured using a Nanodrop (Thermo Fisher Scientific) and evaluated for purity. Samples were treated with DNase (DNA free kit, Life Technologies/Thermo Fisher Scientific) and frozen at -80°C.

### 2.6. RNA Sequencing

RNA samples were sent to Genome Quebec (Montreal, QC) for quality control analyses using an Agilent 2100 Bioanalyzer. RNA integrity was evaluated based on the RNA Integrity Number (RIN), and only samples with a RIN ≥ 7.0 were used for library preparation and sequencing. RNA-seq libraries were prepared using standard Illumina protocols and sequenced on an Illumina platform (NovaSeq PE 100 – 25M reads) at Genome Québec.

Initial quality assessment of raw RNA-seq reads was conducted using FASTQC v0.12.1 [20]. Based on quality metrics, trimming of adapters and low-quality bases was carried out using TRIMMOMATIC software (v0.39;) [21]. The resulting high-quality reads were aligned to the GRCm38 reference genome with STAR v2.7.11b [22], yielding an average unique alignment rate of approximately 90%. Gene-level count matrices were generated using FeatureCounts v2.0.6 with GRCm38 genome annotations (release 101) [23]. Differential gene expression analysis was conducted using the DESeq2 R package [24]. Venn diagram and graphs representing differentially expressed genes (DEGs) were generated based on z-score–transformed normalized counts and performed by R studio (version 2024.04.1+748). Functional enrichment analyses, including Gene Ontology, were performed using the gprofiler2 R package [25] and presented by bubble graph and bar chart performed by R studio. Bioinformatics analyses were performed at the Bioinformatics core facility of the Montreal Clinical Research Institute (Montreal, QC). The raw RNA sequencing data have been submitted to the NCBI BioProject under accession number of PRJNA1422712. Sequence Read Archive (SRA) will be submitted and Permanent accession numbers will be provided upon assignment.

### 2.7. Ingenuity Pathway Analysis

The data derived from RNA sequencing were analyzed to identify enriched canonical pathways, upstream regulators, and associated biological functions and diseases across experimental conditions. DEGs were filtered using |log_2_FC| ≥ 2 and adjusted p< 0.05. The resulting DEGs were imported into Ingenuity Pathway Analysis software (IPA;QIAGEN, Germantown, MA) and Core Analysis performed using the Ingenuity Knowledge Base as the reference dataset, incorporating both direct and indirect molecular interactions. Canonical pathway enrichment was assessed using right-tailed Fisher’s exact test, and pathways with p< 0.05 were considered significantly enriched. Pathway activation or inhibition states were predicted using IPA’s activation *z*-score algorithm. Upstream Regulator Analysis was conducted to identify transcription factors, cytokines, and signaling molecules predicted to explain the observed gene expression changes, with significance determined by overlap *p*-value and activation *z*-score. Additionally, Diseases and Functions analysis was used to associate DEGs with biological functions and disease categories relevant to neuroinflammation and microglial activation, including immune response, inflammatory signaling, and neurological disease processes. Gene interaction networks were generated using IPA’s scoring system, which ranks networks based on the number of focus genes and the connectivity within the Ingenuity Knowledge Base.

### 2.8. Cell culture

Murine microglia BV2 cells kindly provided by Dr. N Sonenberg (Goodman Cancer Research Institute, Department of Biochemistry, McGill University, Montreal, QC) were cultured in DMEM/F12 medium (Wisent, lnc. St-Jean-Baptiste QC), enriched with 10% fetal bovine serum (Sigma-Aldrich, St. Louis, MO), and 1% penicillin/streptomycin (Wisent, lnc.). The cells were incubated at 37 °C with 5% CO_2_ and culture medium was changed with fresh pre-warmed medium every 2–3 days, based on the morphology and confluency of BV2 cells.

### 2.9. In vitro Experimental Protocol

In an effort to elucidate PS-NPs effects, *in* vitro experiments with BV2 cells were conducted. In the first experiment, cells were exposed to different concentrations of PS-NPs for 3, 24, 48, and 72 h, and the respective endpoints analyzed. In the second experiment, BV2 cells were treated with varying concentrations of PS-NPs for 1, 3, 6, 12, 24, 48, and 72 h. Following each exposure period, the treatment medium was replaced with fresh culture medium without PS-NPs (designated as washout period), and the cells were maintained for an additional 48 or 72 h. The selected time points were designed to accurately reflect the temporal pattern of BV2 cell activation and inflammation in relation to PS-NPs internalization dynamics. To avoid confounding effects related to cell confluency, untreated controls cultured at 24, 48, and 72 h were included. This allowed assessment of whether prolonged or residual effects persisted after the removal of PS-NPs.

### 2.10. Cell Viability

BV2 cells were seeded in 96-well plates at a density of 1 × 10⁴ cells/well in 200 µL of complete DMEM/F12 and allowed to adhere overnight. Cells were then exposed to PS-NPs (100 nm or 500 nm) at concentrations of 0, 10, 50, and 100 μg/mL for 3, 24, 48, and 72 h. Following treatment, medium containing PS-NPs was removed and each well washed twice with PBS. A 100 µL aliquot of fresh medium containing 0.5 mg/mL of MTT solution was added to each well and incubated at 37 °C for 3 h. The supernatant was carefully removed, and the resulting formazan crystals were dissolved in 100 µL of DMSO with gentle shaking for 10 min. Absorbance was measured at 570 nm using a MBIZLQ-100 microplate reader (MBI, Montreal Biotech, QC). Cell viability percentage was calculated relative to untreated controls.

### 2.11. PS-NPs internalization assessment

To confirm the internalization of PS-NPs, BV2 cells were seeded on glass cover slips in 24 well plates at a density of 1 × 10^5^ cells/well in 500 µL of complete DMEM/F12 and allowed to adhere overnight. Cells were exposed to fluorescent labeled PS-NPs (100 nm or 500 nm) at concentrations of 10 and 50 μg/mL for 24 h. After exposure, the medium was removed, cells in each well were washed three times with PBS and fixed in 4% paraformaldehyde (PFA) for 10 min at room temperature. The nuclei and plasma membranes were then counterstained with Hoechst dye and Wheat Germ Agglutinin (WGA;1μg/mL; Fisher Scientific) and incubated for 5 min at room temperature in the dark. After staining, cover slips were washed three times with PBS and mounted on glass slides with Fluoromount-G and imaged using a Zeiss LSM 980 laser-scanning confocal microscope. Fluorescence was acquired using standard channels for the labeled PS-NPs (Alexa 488/FITC), WGA conjugate (Alexa 594/TRITC), and nuclear stain (Hoechst).

### 2.12. PS-NPs co-localization with endolysosomal compartments

To investigate the intracellular trafficking and localization dynamics of PS-NPs in BV2 microglia cells, co-localization analyses with endolysosomal markers was done. BV2 cells were seeded on glass coverslips in 24-well plates at a density of 1 × 10⁵ cells/well in 500 µL of complete DMEM/F12 and allowed to adhere overnight. Cells were then exposed to fluorescently labeled PS-NPs (100 nm or 500 nm) at concentrations of 10 µg/mL and 50 µg/mL for 24 h. After exposure, the medium was removed and each well washed three times with PBS, fixed with 4% PFA for 10 min at room temperature. Cells were then washed three times in PBS and permeabilized with 0.1% Triton X-100 for 15 min at room temperature. Following permeabilization, cells were washed and blocked with 5% BSA for 1 h at room. Cells were then washed and incubated overnight at 4°C with rabbit anti-mouse primary antibodies (Cell Signaling Technology, New England Biolabs, Whitby ON) for EEA1 (0.5 µg/mL; early endosome marker), Rab7 (0.5 µg/mL; late endosome marker), LAMP2 (0.2 µg/mL; lysosome marker), and rabbit anti-mouse primary antibody (BioLegend, San Diego, CA, USA) for Giantin (0.2 µg/mL; Golgi apparatus marker). Cells were then washed with PBS and incubated with Alexa Fluor 647-conjugated anti-rabbit secondary antibody (Jackson ImmunoResearch Laboratories) 1 h at room temperature in the dark. Nuclei were counterstained with Hoechst 33258 (1 µg/mL) for 5 min at room temperature, followed by three PBS washes. Coverslips were mounted on glass slides using Fluoromount-G, and images were acquired using a Zeiss LSM 980 laser-scanning confocal microscope.

### 2.13. PS-NPs Uptake kinetics and accumulation

To quantify cellular uptake and examine both uptake kinetics and accumulation of PS-NPs after cessation of exposure, cells were seeded in 24-well plates at a density of 1 × 10^5^ cells/well in 500 µL of complete DMEM/F12 medium and allowed to adhere overnight. Cells were then exposed to fluorescently labeled PS-NPs (100 nm and 500 nm) at concentrations of 10 or 50 μg/mL for 1, 3, 6, 12, 24, and 48 h. After exposure, the medium was removed and each well was washed three times with PBS and fresh clean medium added to each well, and cells were then incubated for up to 48 h. After incubation, cells were washed three times with PBS, fixed in 2% PFA, collected, centrifuged and resuspended in PBS. PS-NPs uptake was quantified using a 4-laser BD LSRFortessa (BD Biosciences, Franklin Lakes, NJ) flow cytometer with 488-nm excitation and 525-nm emission (FITC channel) wavelengths. More than 10,000 events/sample were acquired, data were gated on single, viable cells, and fluorescence compensation was applied using PS-NP-negative controls. Analyses were performed with BD FACSDiva™ (ver 6.2) software.

### 2.14. Microglia activation and inflammation

To investigate the effects of PS-NPs on microglia, two specific markers including Iba1 and CD68 were used. CD68 (Cluster of Differentiation 68) is a type I transmembrane glycoprotein predominantly localized in the lysosomal and endosomal membranes of microglia and functions mainly in phagocytic activity, intracellular digestion, and antigen processing, reflecting the activation and clearance roles of innate immune cells. To assess the expression of these markers, BV2 cells were seeded on glass coverslips in 24-well plates at a density of 1 × 10⁵ cells/well in 500 µL of complete DMEM/F-12 and allowed to adhere overnight. Cells were then exposed to fluorescently labeled PS-NPs (100 nm or 500 nm) at concentrations of either 10 or 50 µg/mL for 1, 3, 6, 12, 24, 48, and 72 h. After exposure, the medium was removed, and each well was washed three times with PBS. Fresh medium was then added to each well, followed by up to 72 h of incubation. After completing the 72h interval, the medium was removed, and each well was washed three times with PBS and fixed with 4% PFA for 10 min at room temperature. After fixation, cells were washed three times in PBS and permeabilized in 0.1% Triton X-100 for 15 min at room temperature. Following permeabilization, cells were washed and blocked with 5% BSA for 1 h at room temperature to minimize nonspecific binding. Subsequently, cells were washed and incubated overnight at 4°C with the following primary antibodies: rabbit anti mouse Iba1 (1 µg/mL; Fujifilm), and rat anti mouse CD68 (2 µg/mL; Thermo Fisher Scientific). Following primary antibody incubation, cells were washed with PBS and then incubated with an Alexa Fluor 647-conjugated anti-rabbit secondary antibody for Iba1 and Alexa Fluor 647-conjugated anti-rat secondary antibody (Jackson ImmunoResearch Laboratories) to detect CD68 for 1 h. Cells were counterstained with Hoechst dye, washed in PBS, mounted with Fluoromount-G, and examined with a Zeiss LSM 980 laser-scanning confocal microscope. To analyze the data, the fluorescent intensity of each cell was measured by MATLAB software (Natick, MA) after precise cell segmentation. Confocal images were first pre-processed to remove background noise and were checked for uneven illumination. Individual cells were identified by nuclear segmentation based on Hoechst staining, followed by cytoplasmic region definition using intensity-based expansion from the nuclear mask. For each segmented cell, mean fluorescence intensity corresponding to either Iba1 or the fluorescent PS-NP signal was extracted. Segmentation accuracy was visually inspected and manually validated to ensure correct cell identification and for exclusion of overlapping or damaged cells. Fluorescence intensity values were quantified per cell, and data from each experimental condition were compared with the untreated control group. The resulting intensity values were used for statistical comparison across nanoparticle size (100 nm vs 500 nm), concentration (10 µg/mL vs 50 µg/mL), and exposure or washout time points.

To explore if the microglia activation is correlated with the accumulation rate of the PS-NPs, the number of PS-NPs particles versus the average fluorescent intensity of Iba1 per each cell was quantified and analyzed by MATLAB software (Natick, MA, USA) using precise cell segmentation in all conditions. Data derived by MATLAB was then analyzed according to followed the Pearson’s correlation coefficient (R) analysis. R studio software was used to present the data.

### 2.15. IL-6 expression analysis

To assess the inflammatory response of BV2 cells following exposure to PS-NPs, cells were seeded in 24-well plates at a density of 1 × 10⁵ cells/well in 500 µL of complete DMEM/F-12 medium and allowed to adhere overnight. Cells were then exposed to fluorescently labeled PS-NPs (100 or 500 nm) at concentrations of either 10 or 50 µg/mL for 3, 24, and 48 h. Lipopolysaccharide (LPS) (MilliporeSigma, Burlington, MA, USA) was included in a separate set of wells as a positive control. Subsequent to PS-NPs exposure, the medium was removed, cells were washed three times with PBS, and fresh medium was added, followed by up to 48 h of incubation. Cells were washed, trypsinized, centrifuged, fixed with 2% PFA, washed, and permeabilized with 0.1% Saponin. Following permeabilization, cells were washed, blocked with 5% BSA, and incubated for 1 h at room temperature with rat anti-mouse IL-6 primary antibody (Biolegend, San Diego, CA, USA). Cells were then washed with PBS and incubated with Alexa Fluor 647-conjugated anti-rat secondary antibody. The IL-6 expression was assessed by fluorescent quantification on a 4-laser BD LSRFortessa (BD Biosciences) using 488-nm excitation/525-nm emission for PS-NPs, and 647-nm excitation/670 emission for Alexa Fluor 647 conjugated antibody. More than 10,000 events per sample were acquired, and data were gated on single, viable cells, and fluorescence compensation was applied using unstained cells, PS-NPs-negative, IL-6-negative, only secondary antibody, only PS-NPs, and TM4 cells (as negative IL-6 expression control) controls. Analyses were performed on the same instrument (BD FACSDiva™ ver 6.2).

### 2.16. Statistical analysis

The data represent the mean of different independent replicate experiments. Analyses were performed using GraphPad Prism 10 software (GraphPad Software Inc., San Diego, CA). One-way and two-way ANOVA analyses were performed, using Dunnett’s multiple comparison test for normally distributed data, or the Kruskal-Wallis test for non-parametric data. Pearson’s correlation coefficient R was used for correlation assessments and statistical significance was defined as *p ≤ 0.05, **p ≤ 0.01, and ***p ≤ 0.001.

## 3. Results

### 3.1. Accumulation of PS-NPs and their effects in cortical region of the mice brain

No significant alterations in body weight, behavior, or general health were noted after 60 days in any of the PS-NP-exposed groups relative to the control group (Fig 1A). Experiments using fluorescently labeled PS-NPs of either 100 or 500 nm were conducted in which mice were administered PS-NPs via daily gavage for 7 days. The presence of fluorescently-labelled PS-NPs of both 100 nm and 500 nm was confirmed by confocal microscopy in the cortex of the brain after 7 days of exposure (Fig 1B). However, morphological analyses of brain sections from all groups, as evaluated by H&E staining, revealed no significant changes in the cortex of the brain for either the 100 nm or 500 nm PS-NPs groups as compared to controls, even after 60 days of PS-NPs administration (Fig 1C).

**Figure 1.**
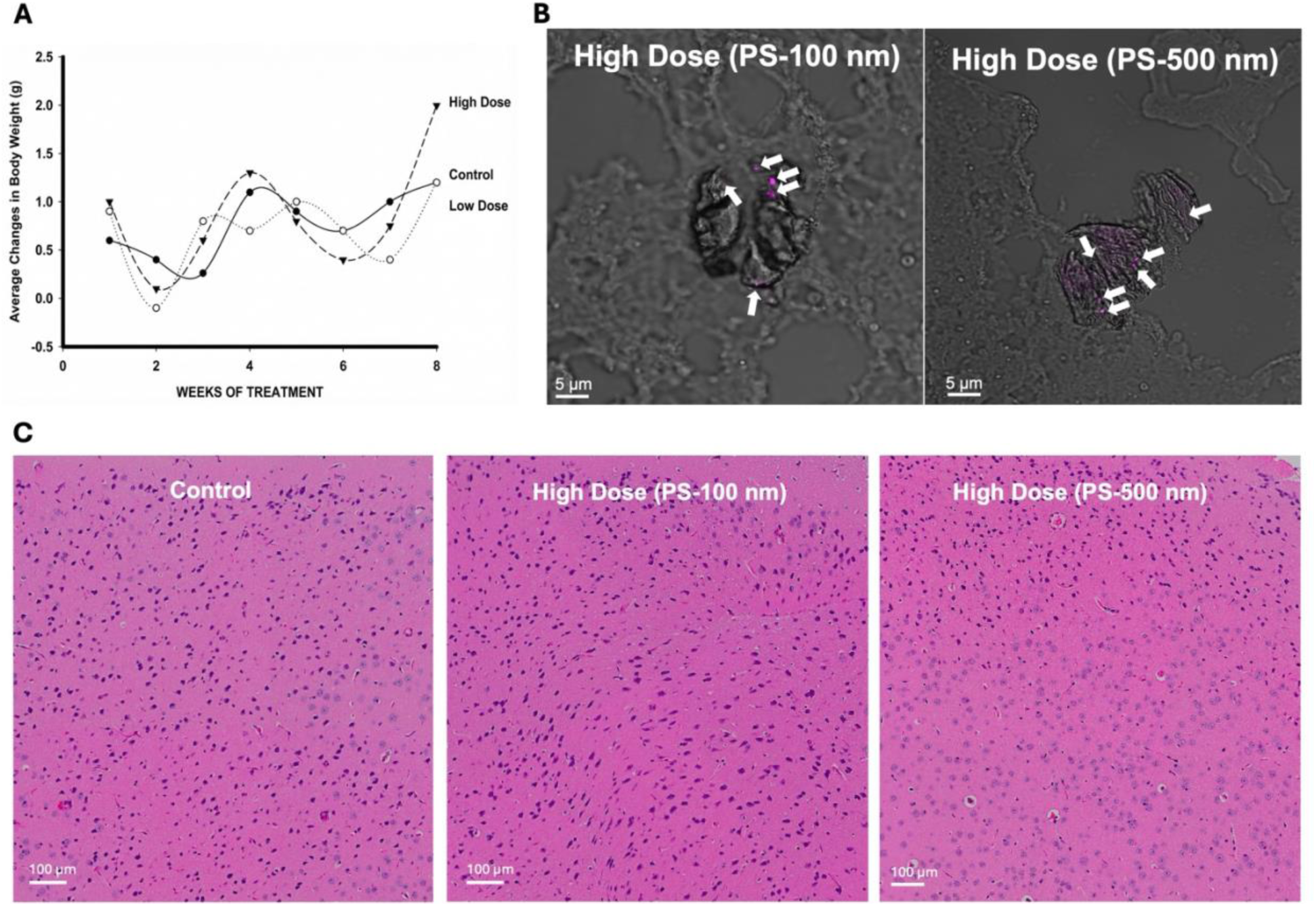
Effects of oral exposure to polystyrene nanoplastics (PS-NPs) on body weight and brain morphology. (A) Relative changes in body weight throughout the study duration. (B) Representative confocal microscopy images showing the presence of fluorescent PS-NPs (100 nm and 500 nm) in the mouse cerebral cortex after 7 days of exposure (arrows indicate internalized particles). (C) Representative hematoxylin and eosin (H&E) sections of brain sections from mice treated for 60 days with PS-NPs (100 and 500 nm). No overt histopathological or gross morphological alterations were observed between treatment groups compared with controls.

### 3.2. Transcriptomics analysis in cortex of mice

Analysis of differentially expressed genes stratified by fold-change magnitude revealed a clear size-dependent transcriptional response in the cortex (Fig 2). Specifically, exposure to 100 nm PS-NPs triggered modest but detectable shifts in gene expression profiles within cortical tissue. Applying the same criteria used in the volcano plots (padj < 0.05 and |log_2_FC| ≥ 2), only 18 genes were significantly altered, including 5 upregulated and 13 downregulated transcripts (Fig 2A). No genes exceeded a 4-fold upregulation threshold, and only a small subset of transcripts showed strong downregulation (≥4-fold). Despite the modest scale of differential expression, the affected genes preferentially map to pathways associated with immune modulation and glial function, suggesting that subtle microglial responses contribute to the observed transcriptional changes rather than widespread tissue disruption. Upstream-regulator analysis revealed several regulators predicted to be activated in the cortex after exposure to 100 nm PS-NPs, including AHR, HNF1A, INSR, and LHX1 (Fig 2B; orange). These regulators were connected to genes linked to protein synthesis and secretory activity (*PRL*), neurotransmitter transport (*SLC18A2, CALCA),* xenobiotic sensing and metabolism (*CYP2E1, CES1F, ALDOB, FBP1*), and early immune/inflammatory signaling (*IL22/IL22R2, CCL5, ITGAD*) among the most prominently connected targets. CREB1 was the only regulator predicted to be inhibited (Fig 2B; blue). Its downstream pattern involved downregulated and moderately upregulated transcripts associated with cellular resilience, vesicle trafficking, and neuronal communication. CREB1 inhibition is consistent with reduced trophic signaling and diminished neuron–glia communication, which are often early features of microglial priming. Diseases and Functions analysis supported these findings. Categories such as behavior, organ morphology, protein synthesis, molecular transport, immune cell trafficking, neurological disease, and nervous system development were significantly enriched. Many of these reflect pathways in which microglia play an essential regulatory role—especially in synaptic maturation, cytokine responsiveness, and neuroimmune communication (Fig 2C).

**Figure 2.**
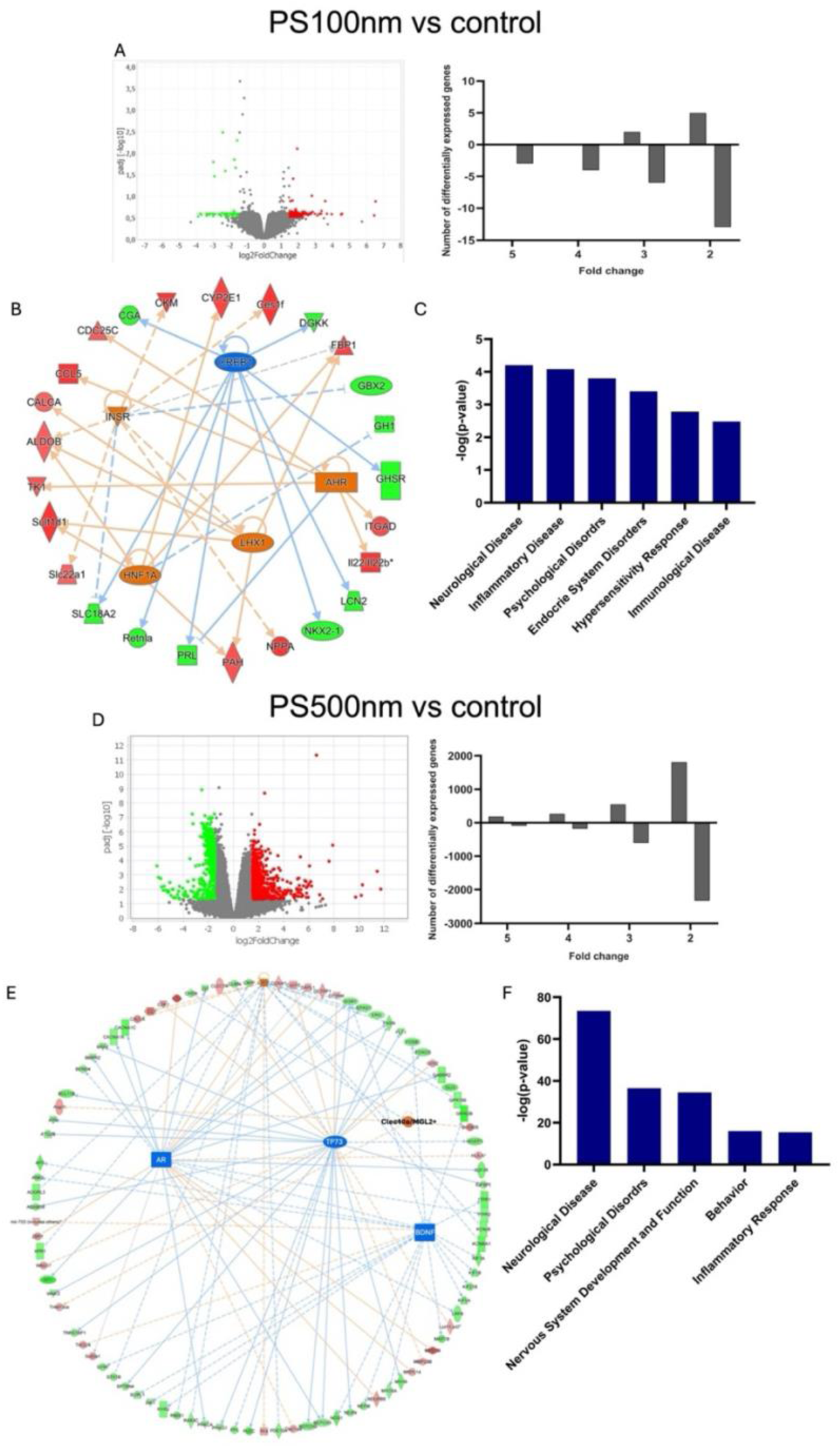
Transcriptomics alterations and upstream regulatory networks induced by exposure to 100 nm PS-NPs. (A) Volcano plots showing up- (red) and down (green) regulated genes. (B) IPA analysis of upstream regulatory network illustrating predicted activation (orange) and inhibition (blue) of key regulators. Colored arrows correspond to activation (orange), and inhibition (blue) regulatory direction based on curated IPA knowledge as Solid (direct) or dotted (indirect) lines. (C) Top enriched biological pathways in disease and functions identified by IPA (−log_10_ *p-*value). (D) Transcriptomics alterations and upstream regulatory networks induced by exposure to 500 nm PS-NPs. Volcano plot showing upregulated (red) and downregulated (green) genes. The number of significantly up- and down-regulated genes displayed according to fold-change magnitude following PS-NP exposure. Differentially expressed genes were identified using a significance threshold of p < 0.05 and absolute |log_2_FC| ≥ 2, ≥ 3, ≥ 4, and ≥ 5. (E) IPA-derived upstream regulator network illustrating predicted activations (orange) and inhibitions (blue) driving the observed transcriptional profile. (F) Top enriched biological pathways involved in disease and functions (−log_10_ *p*-value) identified by IPA.

In contrast, more extensive alterations in gene expression were observed in cortical samples from the 500 nm PS-NPs group as compared with controls, indicating size-dependent transcriptional effects. Differential expression analysis identified 8,954 significantly altered genes (4,146 upregulated and 4,808 downregulated; padj < 0.05), reflecting a large-scale perturbation of gene expression (Fig 2D). When applying a biologically stringent cutoff (padj < 0.05 and |log_2_FC| ≥ 2), 4,148 genes remained differentially expressed, including 1,815 upregulated and 2,333 downregulated transcripts. A substantial fraction of these genes displayed high changes in expression, with 548 genes exceeding 3-fold change, 265 exceeding 4-fold change, and 186 exceeding 5-fold change, underscoring the magnitude of transcriptional remodeling induced by larger particles. Many of the most responsive genes are associated with microglial activation, cytoskeletal remodeling, inflammatory signaling, and immune regulation, consistent with a robust neuroimmune response to 500 nm PS-NP exposure. Upstream-regulator analysis identified TP73 (tumor protein p73), AR (androgen receptor), and BDNF (brain-derived neurotrophic factor) as major predicted inhibited regulators (Fig 2E; blue) and CLEC10A/MGL2 (Macrophage Galactose-type Lectin) as major predicted activated regulator (Fig 2E; orange). Their predicted inhibition and activation was linked to several strongly altered genes associated with inflammatory signaling, structural remodeling, and loss of trophic support. Upregulated downstream targets included innate immune and microglia-associated inflammatory genes such as (*ZBP1, CCL5, CXCL10, IL8, LYZ1/LYZ2)*, together with remodeling-related changes (*MMP28, TAGLN, NOTCH1/NOTCH3*), consistent with a shift toward neuroinflammatory activation occurring alongside predicted TP73/BDNF inhibition and disrupted neuronal function, maintenance, development, and plasticity mechanisms (*SYN1, STX1B, SPTBN4, SORL1, MAP1B, GRIN2B, ITPR1/2, CDH9*). Diseases and Functions analysis supported these findings. Categories such as neurological disease, pcychological disorders, nervous system development and function, behavior, and inflammatory response were significantly enriched (Fig 2F).

The upstream-regulator analysis between 500 and 100 nm PS-NPs exposed groups predicted inhibition of several genes, including TGFB1 (transforming growth factor beta 1) a multifunctional cytokine that regulates immune responses, inflammation, and tissue homeostasis, BDNF (brain-derived neurotrophic factor) a neurotrophins essential for neuronal survival, synaptic plasticity, and neurogenesis, CX3CR1 (C-X3-C motif chemokine receptor 1) a chemokine receptor predominantly expressed by microglia that mediates neuron–microglia communication and immune surveillance in the central nervous system, and F2 (coagulation factor II, thrombin) a serine protease involved in blood coagulation which also modulates inflammation and neurovascular signaling (Fig 3A; blue). These molecules collectively regulate microglial homeostasis, neuroimmune communication, trophic signaling, and chemokine-mediated microglial positioning, as reflected by altered expression of genes involved in immune signaling and cell–cell interactions, including (*CCL5, ITGAD, LCN2, IL6R, HEG1, HIVEP1)*. Their predicted inhibition is therefore consistent with coordinated downregulation of pathways governing neuron–microglia communication, synaptic and ion channel signaling (*GRIN2B, GABBR2, ITPR1/2, KCNQ2/3, KCNJ6*), and tissue remodeling, suggesting that larger particles induce a greater disruption of neurotrophic and regulatory signaling in the cortex. Conversely, CNTF and APOE were predicted to be activated (Fig. 3A; orange). Activation of CNTF- and APOE-associated regulatory networks coincided with altered expression of glial, immune, and metabolic response genes, including (*LCN2, CCL5, ITGAD, PPARA*) and cytoskeletal remodeling-associated genes (*TAGLN, MYH9, MYO5A*), consistent with enhanced glial reactivity and metabolically adaptive microglial states. Comparison between 500 nm PS-NPs and 100 nm PS-NPs exposed groups shared 44 genes with the 500 nm PS-NPs vs control dataset, whereas only 12 genes overlapped with the 100 nm PS-NPs vs control comparison. Notably, 500 nm PS-NPs vs control exhibited 2 unique genes, while 100 nm PS-NPs vs control displayed 5 unique genes, underscoring distinct size-dependent transcriptional signatures. A single gene (*POMC*) was common to all three comparisons (Fig 3B). Magnitude of transcriptional alterations involved in synaptic transmission and plasticity (*GRIN2B, SYN1, STX1B*), neuronal structural organization (*MAP1B, SPTBN4, SHANK2*), junctional and ATP-dependent signaling pathways (*ITPR1, ITPR2*), and neuroendocrine signaling pathway (*POMC*) contributed to the observed IPA analysis network, with more pronounced transcriptional changes detected following exposure to 500 nm PS-NPs. In parallel, immune- and microglia-associated genes (*CCL5, CXCL10, LCN2, LYZ2*) were upregulated, indicating activation of neuroimmune-related biological processes (Fig 3C).

**Figure 3.**
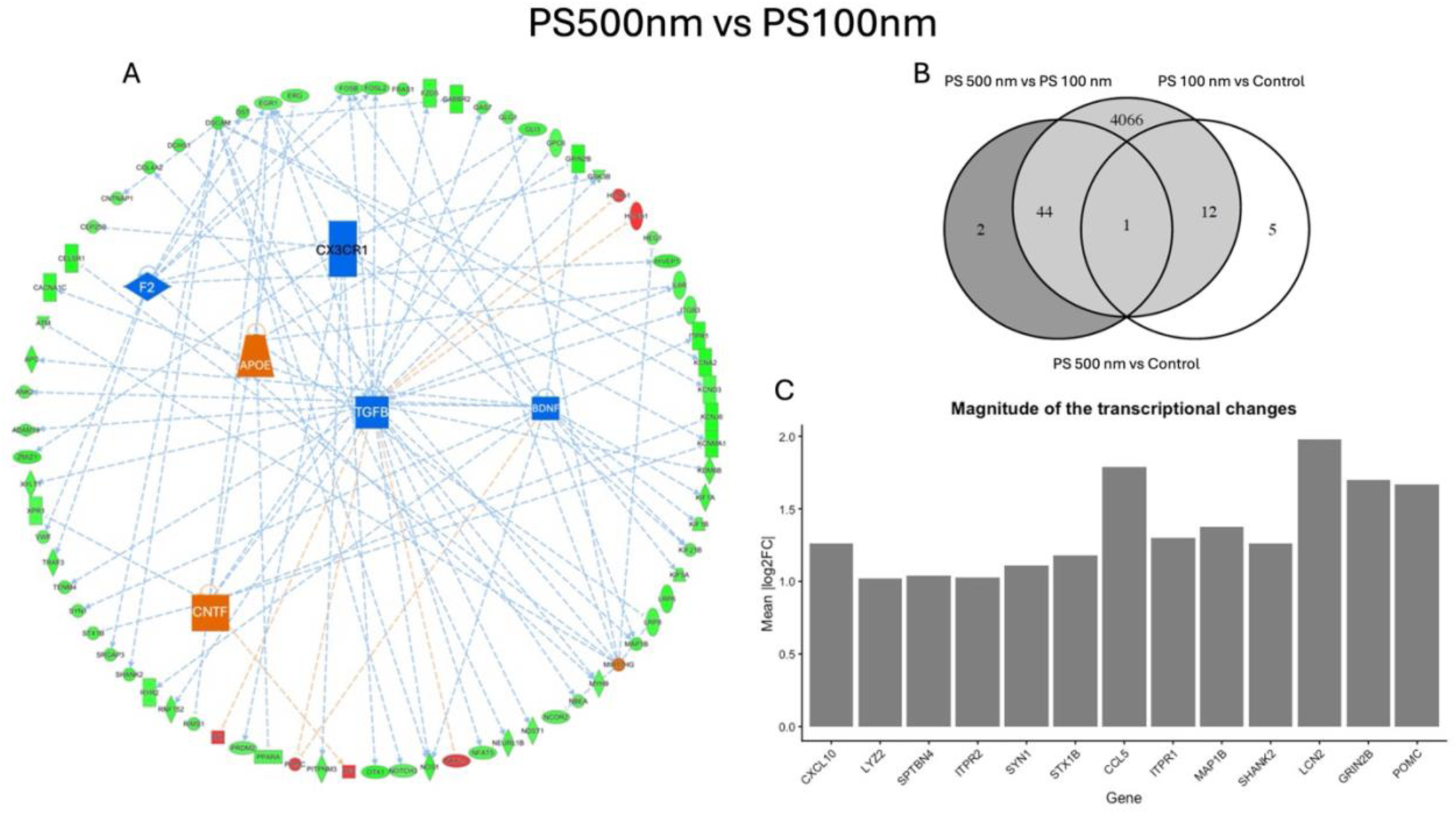
(A) IPA upstream regulator network showing predicted activation (orange) and inhibition (blue) of key regulators orchestrating the transcriptional response between the 100 and 500 nm PS-NPs exposed groups. (B) Venn diagram showing the overlap of differentially expressed genes (DEGs) between controla, 100 and 500 nm exposed groups. among PS-NPs 100 versus Control, PS-NPs 500 versus Control, and PS-NPs 500 versus PS-NPs 100 comparisons. DEGs were defined using the criteria |log_2_FC| ≥ 1 and adjusted *p* value < 0.05.-specific, indicating distinct transcriptional responses driven by nanoplastic particle size. (C) Magnitude of transcriptional changes. Bars show the mean absolute log_2_ fold change (|log_2_FC| ≥ 2) across exposure contrasts for selected genes contributing to enriched IPA network, highlighting strong perturbations in immune/microglia-associated genes, and neuropeptide/neuroendocrine signaling gene (*POMC*).

Using Gene Ontology (GO) Biological Process categories revealed size-dependent transcriptional alterations in the cortex. GO enrichment analysis identified significant modulation of biological processes related to neuroinflammation and immune system activation, regulation of nervous system development, synaptic signaling and neuronal communication, cell junction organization, and cellular stress and metabolic processes (Fig S1).

### 3.3. Effects on Glial Cells

To assess whether chronic PS-NP exposure alters glial cell populations, Iba1–positive microglia and GFAP-positive astrocytes in the cortex were quantified using Skeleton and Sholl [26] (Fig 4). In the vehicle control group, cortical microglia displayed a branched, morphology characterized by long, thin processes extending from a small soma (Fig 4A). Following PS-NPs exposure, microglia adopted a less complex, reactive-like morphology with shortened and simplified branches. Quantitatively, skeleton analysis showed average marked reductions in the number of branches per cell (Fig 4B), maximal branch length (Fig 4C), number of junctions (Fig 4D), and junction voxels (Fig 4E), slab voxels (Fig 4F) and end-point voxels (Fig 4G) in both PS-NP groups compared with vehicle. Sholl analysis revealed a significant decrease in the number of process intersections with concentric circles around the soma (Fig 4I), confirming a collapse of microglial process complexity. Despite these pronounced morphological changes, the overall density of Iba1–positive microglia in the cortex was not significantly altered (Fig 4J), indicating that PS-NPs primarily remodel microglial branching rather than inducing overt microglial loss at this time point.

**Figure 4:**
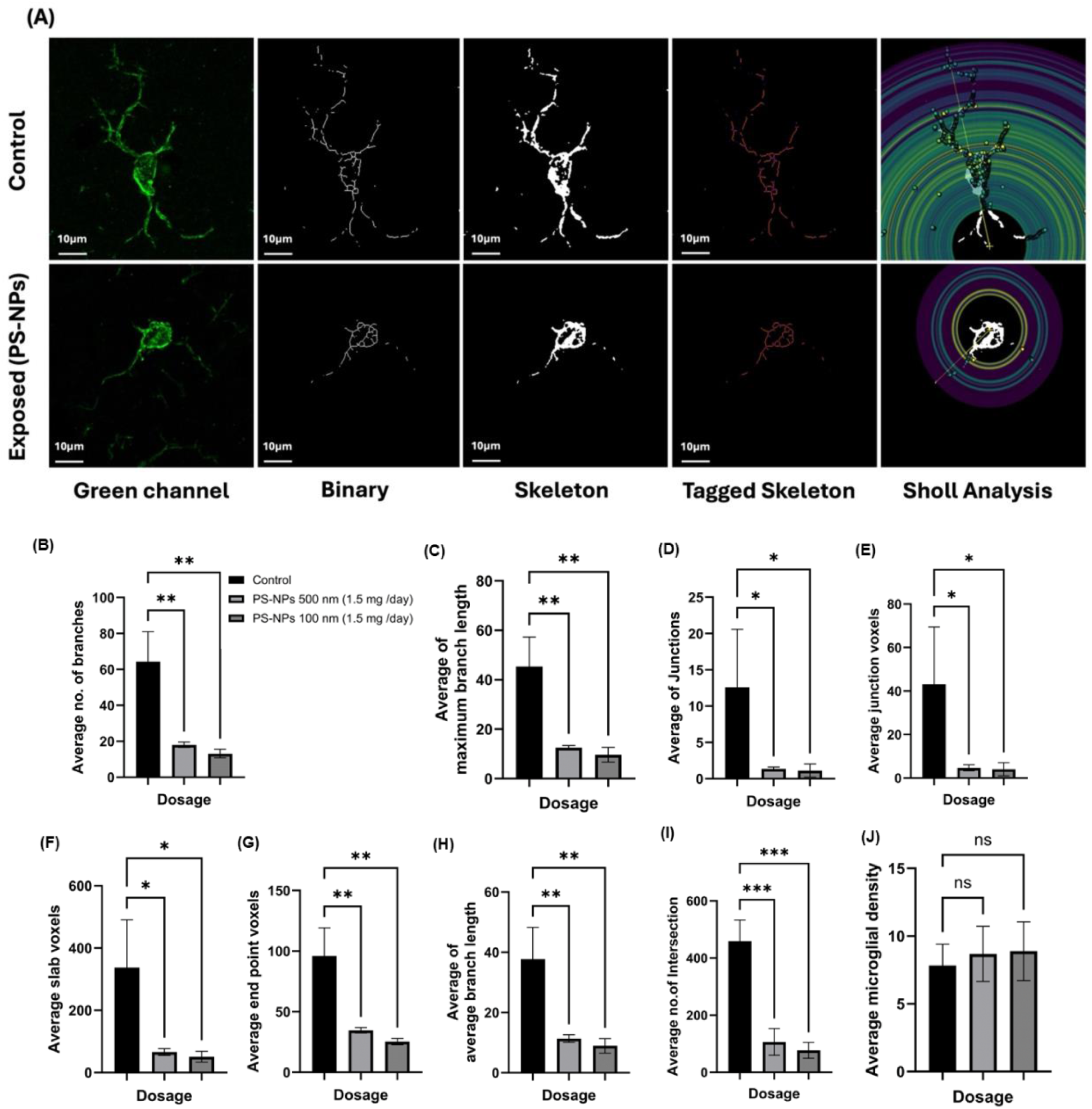
Mice received vehicle (Control), PS-NPs 500 nm (1.5 mg/day/kg body weight) and PS-NPs 100 nm (1.5 mg/day/kg body weight), and microglial morphology in the cortex was quantified by 3-D skeleton analysis of Iba1 immunofluorescence. (A) Representative images show the skeleton and Sholl analysis of cortical microglia in control and PS-NP–exposed mice. The top row shows microglia from control animals, and the bottom row shows microglia after PS-NPs exposure. “Binary” panels depict binarized Iba1–positive microglial processes used as input for quantitative analysis. “Skeleton” panels show 1-pixel-wide skeletons of each cell’s arbor. “Tagged skeleton” panels display the Analyze Skeleton output with branches, junctions, and endpoints pseudo-colored for extraction of morphometric parameters (branch length, numbers of junctions and endpoints, etc.). “Sholl analysis” panels illustrate Sholl analysis, with concentric circles centered on the soma; intersections between processes and each radius were used to quantify process complexity and ramification. All images were processed in FIJI/ImageJ using the Skeletonize, Analyze Skeleton, and Sholl Analysis plugins. Scale bar: 10 µm. Figure 2B onwards: Skeleton analysis of Iba1–positive microglial cells in the cortical region following exposure to 100 nm and 500 nm PS-NPs. (B) Average number of branches per microglial cell. (C) Average maximum branch length. (D) Average number of junctions. (E) Average junction voxels. (F) Average slab voxels. (G) Average end-point voxels. (H) Average branch length. Figure 2I onwards; Sholl analysis of Iba1–positive microglial cells in the cortical region following exposure to 100 nm and 500 nm PS-NPs. (I) Average number of intersections between microglial processes and concentric Sholl circles. (J) Average microglial density. Data are presented as mean ± SD. Statistical significance was assessed by one-way ANOVA followed by Dunnett’s multiple-comparisons test versus control, using multiplicity-adjusted P values. ns, not significant; *P ≤ 0.05, **P ≤ 0.01, ***P ≤ 0.001.

Astrocytes exhibited a comparatively modest structural response to PS-NP exposure. In control conditions, GFAP-positive astrocytes maintained their typical stellate morphology, characterized by multiple fine processes. However, chronic exposure to 100 nm and 500 nm PS-NPs led to a reduction in the average number of GFAP-positive branches per astrocyte, accompanied by significant or near-significant decreases in junction-related parameters and end-point voxels, particularly in the 500 nm group (Fig 5A–H).

**Figure 5:**
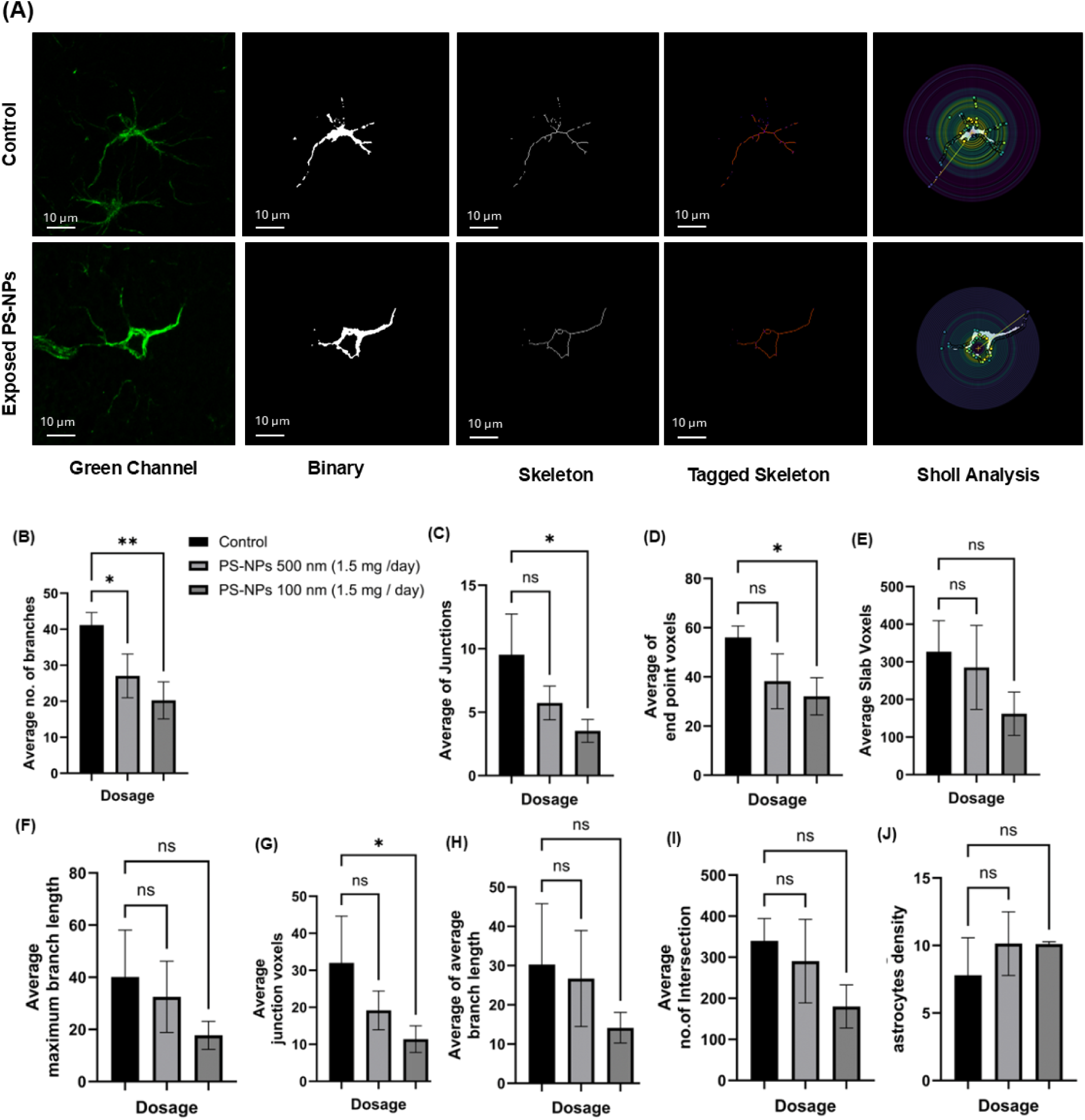
Mice received vehicle (Control), PS-NPs 500 nm (1.5 mg/day/body weight) and PS-NPs 100 nm (1.5 mg/day/body weight), and astrocytes morphology in the cortex was quantified by 3-D skeleton analysis of GFAP immunofluorescence. (A) Representative images show the skeleton and Sholl analysis of cortical astrocytes in control and PS-NP–exposed mice. The top row shows astrocytes from control animals, and the bottom row shows astrocytes after PS-NPs exposure. “Binary” panels depict binarized GFAP –positive astrocytes processes used as input for quantitative analysis. “Skeleton” panels show 1-pixel-wide skeletons of each cell’s arbor. “Tagged skeleton” panels display the Analyze Skeleton output with branches, junctions, and endpoints pseudo-colored for extraction of morphometric parameters (branch length, numbers of junctions and endpoints, etc.). “Sholl analysis” panels illustrate Sholl analysis, with concentric circles centered on the soma; intersections between processes and each radius were used to quantify process complexity and ramification. All images were processed in FIJI/ImageJ using the Skeletonize, Analyze Skeleton, and Sholl Analysis plugins. Scale bar: 10 µm. Skeleton analysis of GFAP–positive astrocytes cells in the cortical region following exposure to 100 nm and 500 nm PS-NPs. Skeleton analysis of GFAP–positive astrocytes cells in the cortical region following exposure to 100 nm and 500 nm PS-NPs. (B) Average number of branches per astrocyte. (C) Average number of junctions. (D) Average end-point voxels. (E) Average slab voxels. (F) Average maximum branch length. (G) Average junction voxels. (H) Average branch length. Figure I onwards; Sholl analysis of GFAP–positive astrocytes cells in the cortical region following exposure to 100 nm and 500 nm PS-NPs. (I) Sholl analysis of GFAP–positive astrocytes cells in the cortical region following exposure to 100 nm and 500 nm PS-NPs. (H) Average number of Sholl intersections per astrocyte. (J) Average density of GFAP-positive astrocytes. Data are shown as mean ± SD and were analyzed by one-way ANOVA followed by Dunnett’s multiple-comparisons test versus control. ns, not significant; *P ≤ 0.05, **P ≤ 0.01, ***P ≤ 0.001.

Quantitative skeleton analysis revealed marked reductions in several structural features, including the average number of branches per cell (Fig 5B), number of junctions (Fig 5C), end-point voxels (Fig 5D), slab voxels (Fig 5E), maximal branch length (Fig 5F), average junction voxels (Fig 5G), average branch length (Fig 5H), and in both PS-NP-treated groups compared with the vehicle controls. Sholl analysis further demonstrated a significant decrease in process intersections with concentric circles around the soma (Fig 5I), confirming a loss of process complexity. Despite these pronounced morphological alterations, the overall density of GFAP-positive astrocytes in the cortex remained unchanged (Fig 5J), suggesting that PS-NPs primarily affect astrocytic branching and morphology rather than cell survival. Collectively, these findings indicate that chronic PS-NP exposure leads to a clear simplification of cortical microglial branching and more subtle remodeling of astrocytic processes, without substantial alterations in glial cell density.

### 3.4. *In vitro* Cell viability

The viability of BV2 cells was altered depending on time/concentration of exposure and size of PS-NPs. A significant decrease in cell viability was observed at 3 h in cells exposed to PS 500 nm, while viability of cells exposed for longer times or cells treated with PS 100 nm remained unchanged. Furthermore, when exposure lasted for 24 h, the viability of BV2 cells remained mostly unchanged. This trend was also observed at 48 and 72 h. (Fig 6).

**Figure 6.**
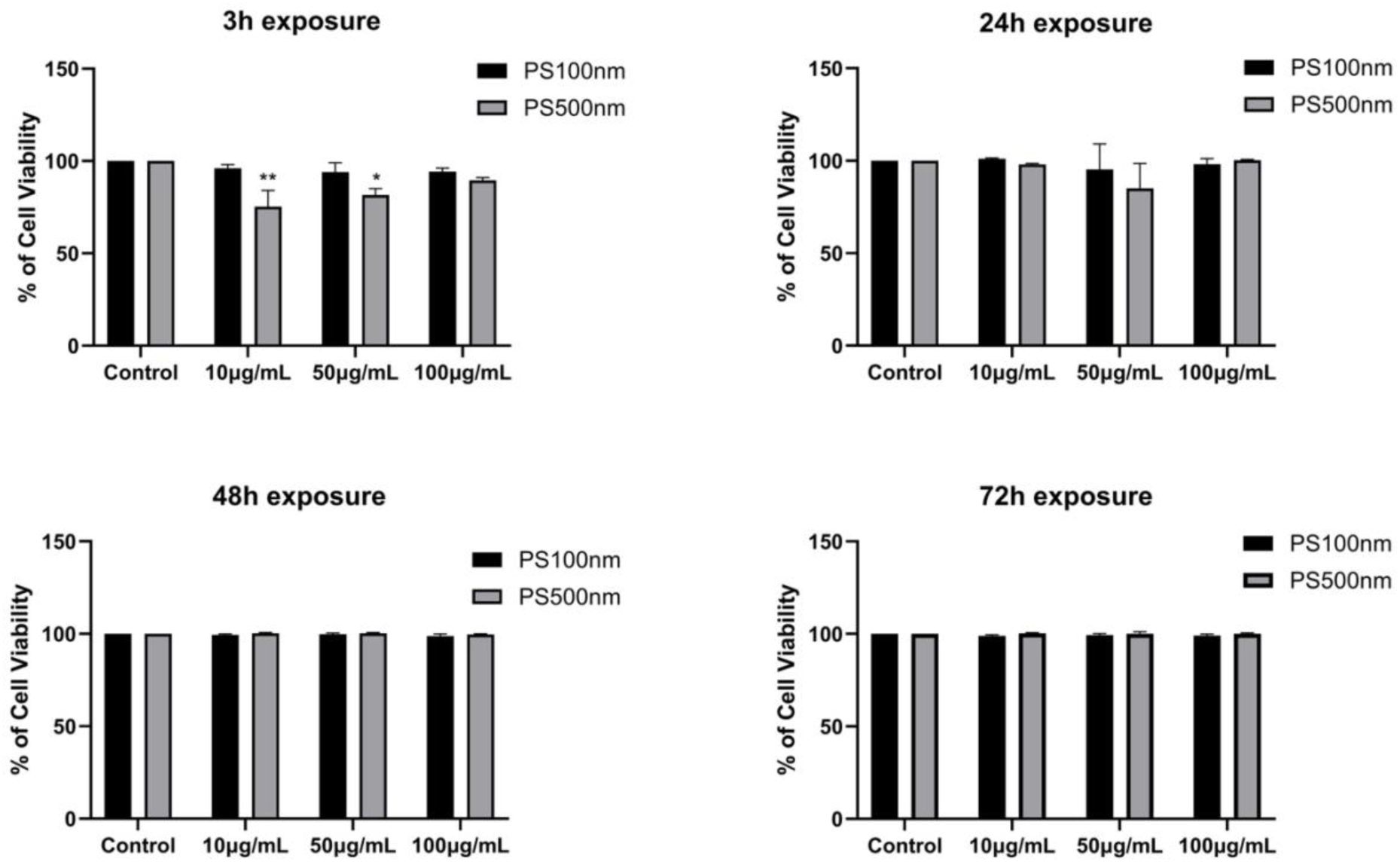
Percentage of viable BV2 cells relative to untreated control after exposure to PS-NPs in sizes of 100 nm and 500 nm at concentrations of 10–100µg/mL for 3–72h of exposure. Data are represented as Mean ± SEM. Two-way ANOVA with Dunnett’s post-test were used for statistical analysis. The statistical significance is indicated in the graph as *P ≤ 0.05, **P ≤ 0.01.

### 3.5. PS-NPs internalization, uptake kinetics, and co-localization

Results showed that BV2 cells were able to internalize the nanoplastics at all exposure concentrations of PS-NPs for 24 h. Confocal microscopy images confirmed that both PS 100 nm and PS 500 nm were present in the cells after 24 h at varying concentrations (Fig 7A).

**Figure 7.**
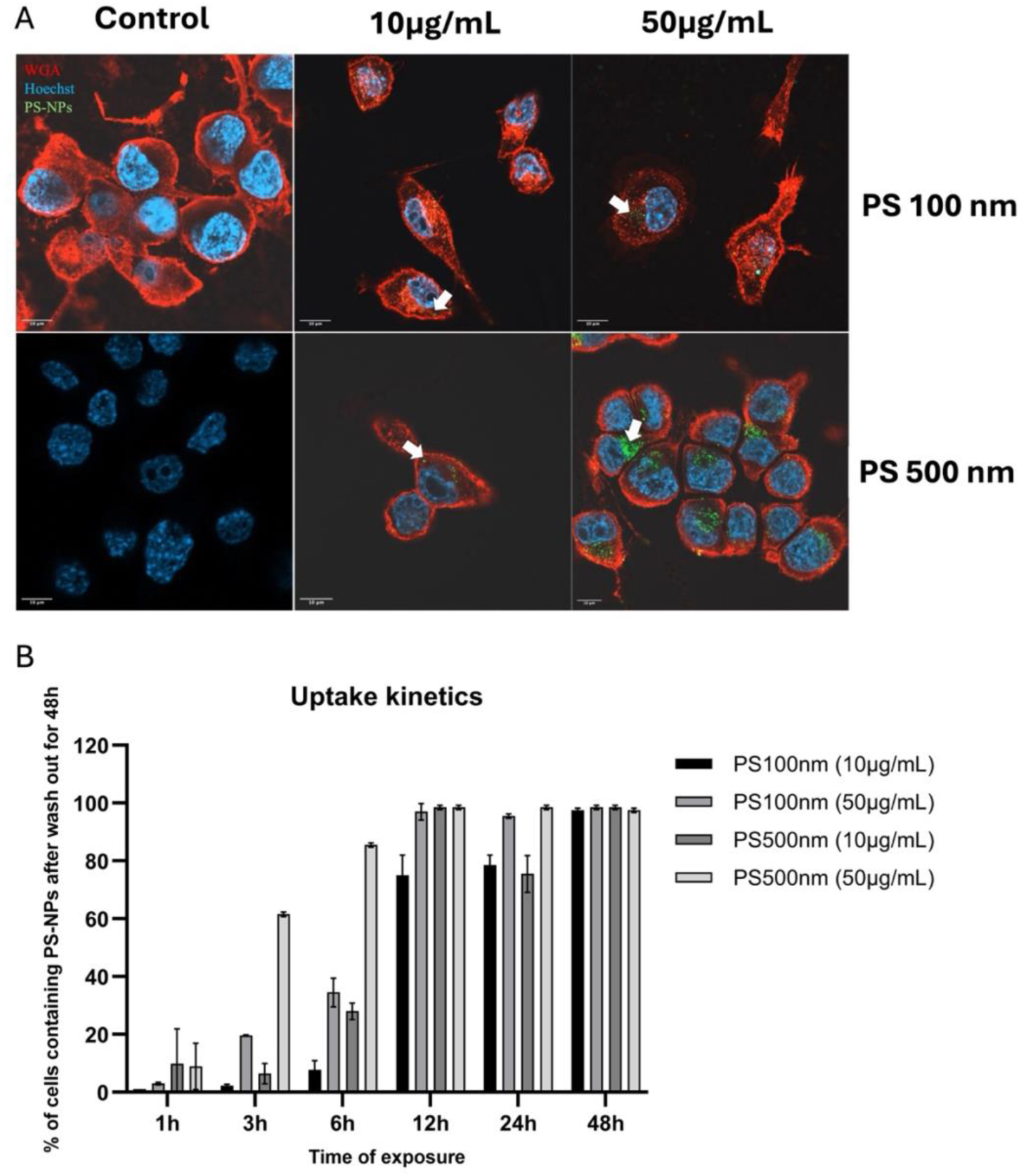
(A) Confocal images showing the internalization of PS-NPs of 100 nm and 500 nm in varying concentrations. The PS-NPs are shown in green, nuclei are in blue, and cell membrane in red. The images are taken from randomly selected areas using the PS-NPs-free sample as negative control. (B) Uptake kinetics of PS-NPs in BV2 cells measured and demonstrated as percentage of cells containing PS-NPs after treatment washout and replacement with PS-NPs free medium. Data are represented as mean ± SD.

Additionally, cellular uptake of PS-NPs was quantified to assess uptake kinetics and accumulation in microglial cells. After exposure/washout conditions, flow cytometry data showed that accumulation of PS-NPs in BV2 cells was dependent on time of exposure and particle size. Approximately 100% of cells displayed PS-NPs accumulation in longer exposure times (12 to 48 h). Size and concentration of PS-NPs affected this accumulation as uptake was minimal at early time points (1–3 h) but increased markedly by 6 h. By 12 h, the majority of cells (>90%) contained PS-NPs, particularly at 50 µg/mL, and uptake reached a plateau by 24 h. At 48 h, near-complete uptake (>95%) was observed across all conditions, indicating efficient and persistent cellular retention of PS-NPs following wash-out (Fig 7B).

To understand the dynamics of internalization, we examined the co-localization of PS-NPs with subcellular compartments. The results show that both sizes of PS-NPs co-localized to, late endosome (Rab7), while 100nm PS-NPs co-localized to lysosome (LAMP2) suggesting that the PS-NPs in BV2 cells follow the endolysosomal pathway (Fig 8A). Quantitative analysis indicated a high correlation between the fluorescence intensity of PS-NPs and both Rab7 and LAMP2 demonstrating that PS-NPs accumulated within the late endosomes and lysosomes (Fig 8B).

**Figure 8.**
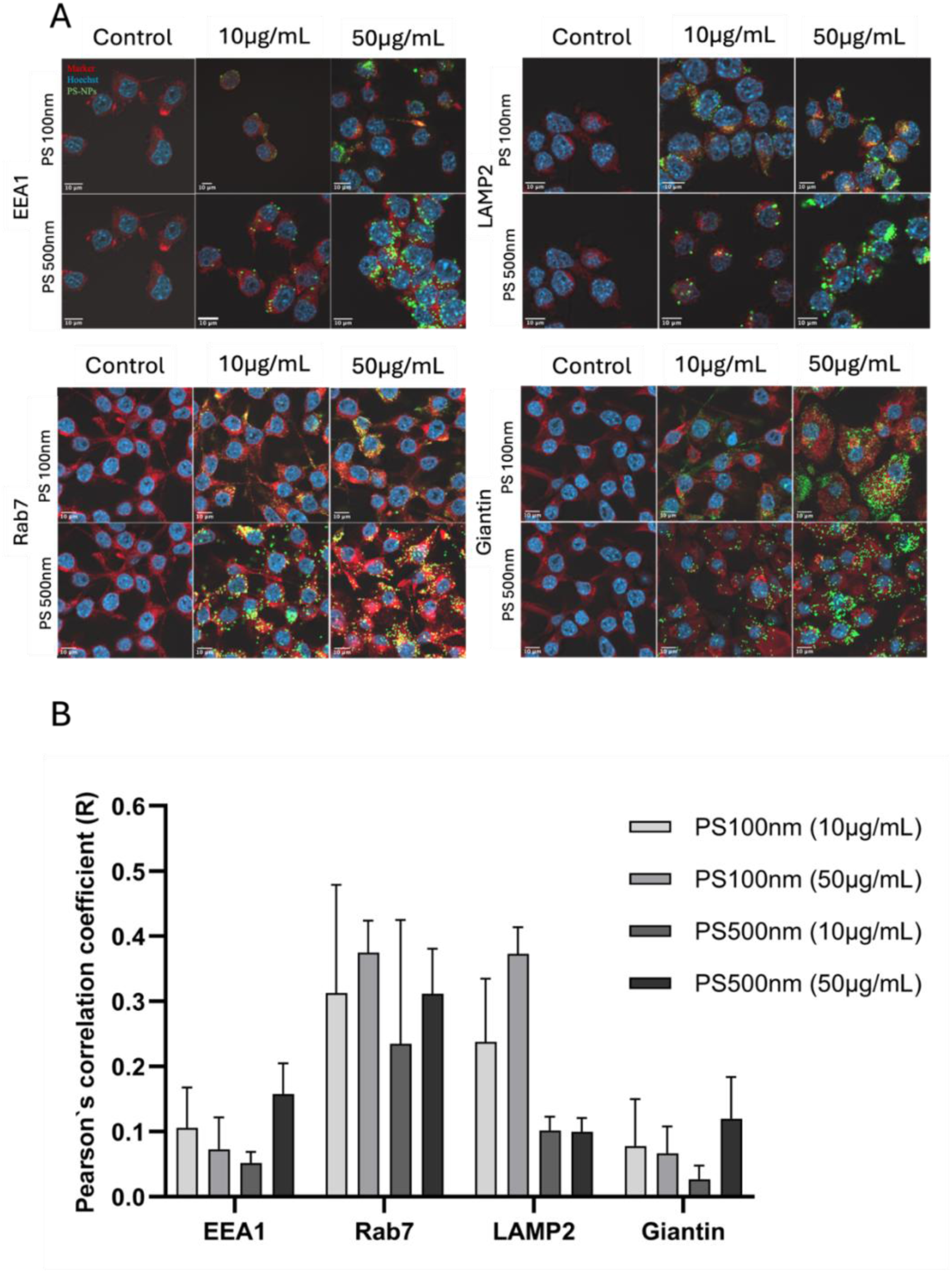
(A) Confocal images representing the co-localization of PS-NPs within endolysosomal compartments. The micrographs are derived from randomly selected areas of immunofluorescent stained cells after appropriate conditions of exposure, representing cell nuclei in blue, PS-NPs in green and specific markers in red; EEA1, Rab7, LAMP2, and Giantin corresponding to early endosome, late endosome, lysosome, and Golgi apparatus, respectively. (B) The correlation of PS-NPs at various concentrations with different compartments of the endolysosomal pathway, as indicated by compartment-specific markers. The data are derived from randomly selected areas by confocal microscopy followed by Pearson’s correlation coefficient (R) analysis quantifying the degree of linear correlation between fluorescent intensities of green channel (PS-NPs) and far-red channel (different markers) within defined regions of interest; average number of 100 cells per condition were considered. Data are represented as R value ± SD.

### 3.6. Activation of Microglia

We next investigated the activation of microglia, which showed transient increases in Iba1 (Fig 9) and CD68 (Fig 10) expression at various times of exposures. Iba1 levels were significantly increased in the PS 100 nm exposed cells at 6, 12, and 24 h of exposure; while in PS 500 nm exposed cells, Iba1 expression increased at 3, 6, 24, and 48 h of exposure, relative to untreated controls (Fig 9A, B). CD68 levels also showed transient changes, in which levels in the low dose 100 nm group peaked at 12 h and in the high dose group peaked at 24 h of exposure (Fig 10A, B). In the cells exposed to PS 500 nm, no significant changes in CD68 levels were observed (Fig 10A, B).

**Figure 9.**
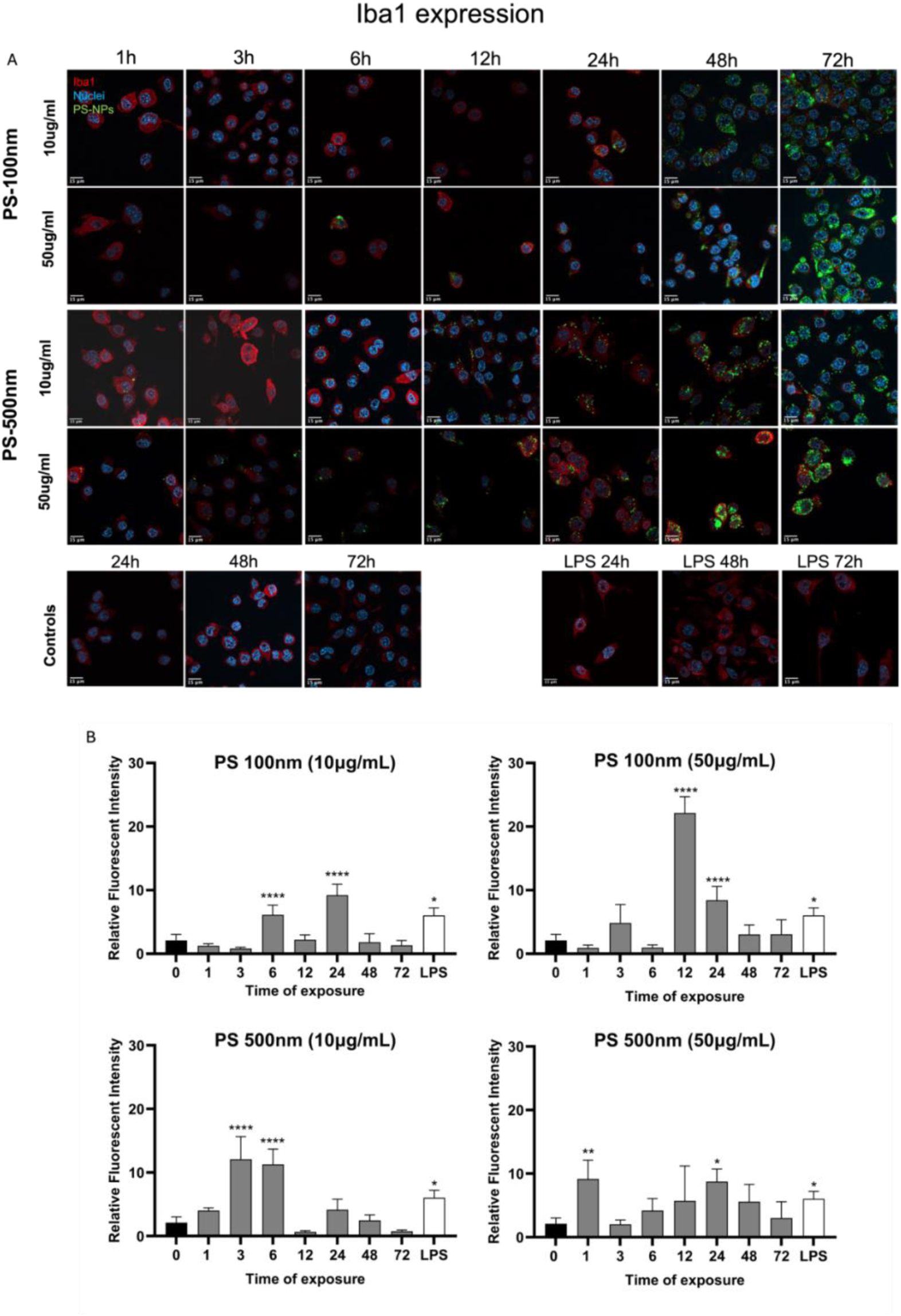
(A) Images of confocal microscopy representing Iba1 expression in BV2 cells at various conditions of exposure. The images were captured from randomly selected areas in each sample and images were obtained at the same magnification in different samples and conditions. Untreated cells were considered as negative controls, and LPS-treated cells were used as positive control for Iba1expression. Nuclei are shown in blue, PS-NPs in green, and Iba1 marker is shown in red. (B) The graph represents average fluorescent intensity of Iba1. As depicted in this graph, microglia cells exhibited transient activation, reflected as Iba1 expression, in response to PS-NPs exposure. Shorter duration of exposure resulted in higher Iba1 expression, in comparison to the longer duration of exposure. Data are presented as Mean ± SEM. One-way ANOVA with Dunnett’s post-test were used for statistical analysis. Statistical significance is indicated as *P ≤ 0.05, **P ≤ 0.01.

**Figure 10.**
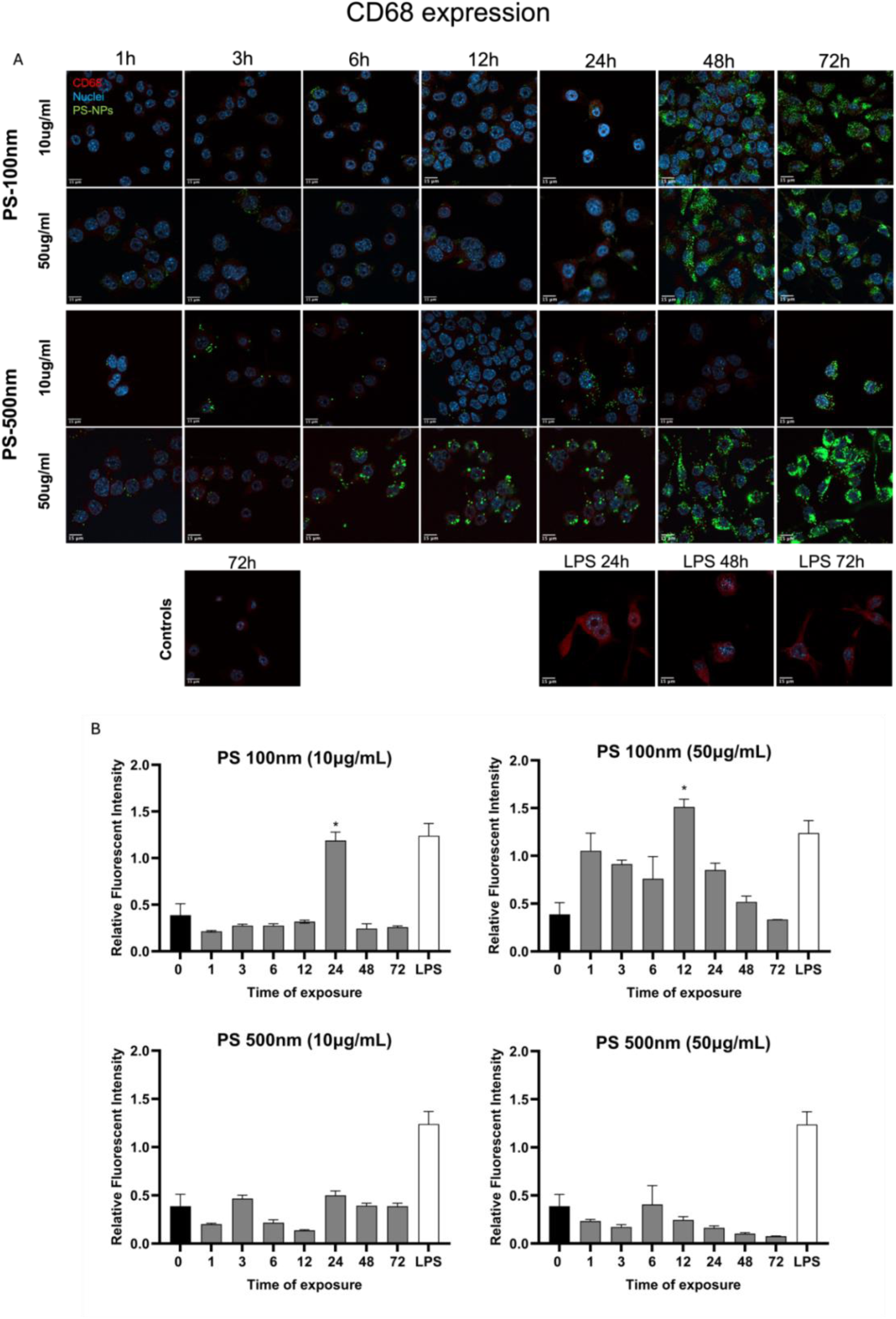
(A) The images of confocal microscopy representing the CD68 expression in BV2 cells at various conditions of exposure. The images were captured from randomly selected areas in each sample and images acquired at identical magnification in different samples and conditions. LPS was used as positive control at various times of exposure. In these images, the nuclei defined as blue, PS-NPs in green and the CD68 marker in red. (B) The graph represents average fluorescent intensity of CD68. As depicted in this graph, microglia cells exhibited transient activation in response to PS-NPs exposure and in the shorter times of exposure the CD68 expression is higher in comparison to the longer times of exposure. Data are presented as the Mean ± SEM. One-way ANOVA with Dunnett’s post-test were used for statistical analysis. The statistical significance is indicated in the graph as *P ≤ 0.05.

### 3.7. Correlation between PS-NPs accumulation and microglial activation

Single cell analysis revealed a time- and concentration-dependent relationship between the number of PS-NPs and microglial activation. Weak or negligible correlations were observed at early time points (1–3 h) and at lower concentrations, indicating limited particle internalization and Iba1 expression during initial exposure. In contrast, positive correlations emerged at 6 h (10 µg/mL) and 24 h (50 µg/mL). At the later time points (48–72 h), correlation coefficients decreased, consistent with the transient activation profile observed in Iba1 and CD68 expression analyses (Fig 11). These data indicate that microglial activation is not linearly proportional to cumulative PS-NP uptake but instead follows a dynamic and transient relationship, supporting a model in which microglial responses are driven by intracellular particle sensing and adaptive regulation rather than continuous accumulation.

**Figure 11.**
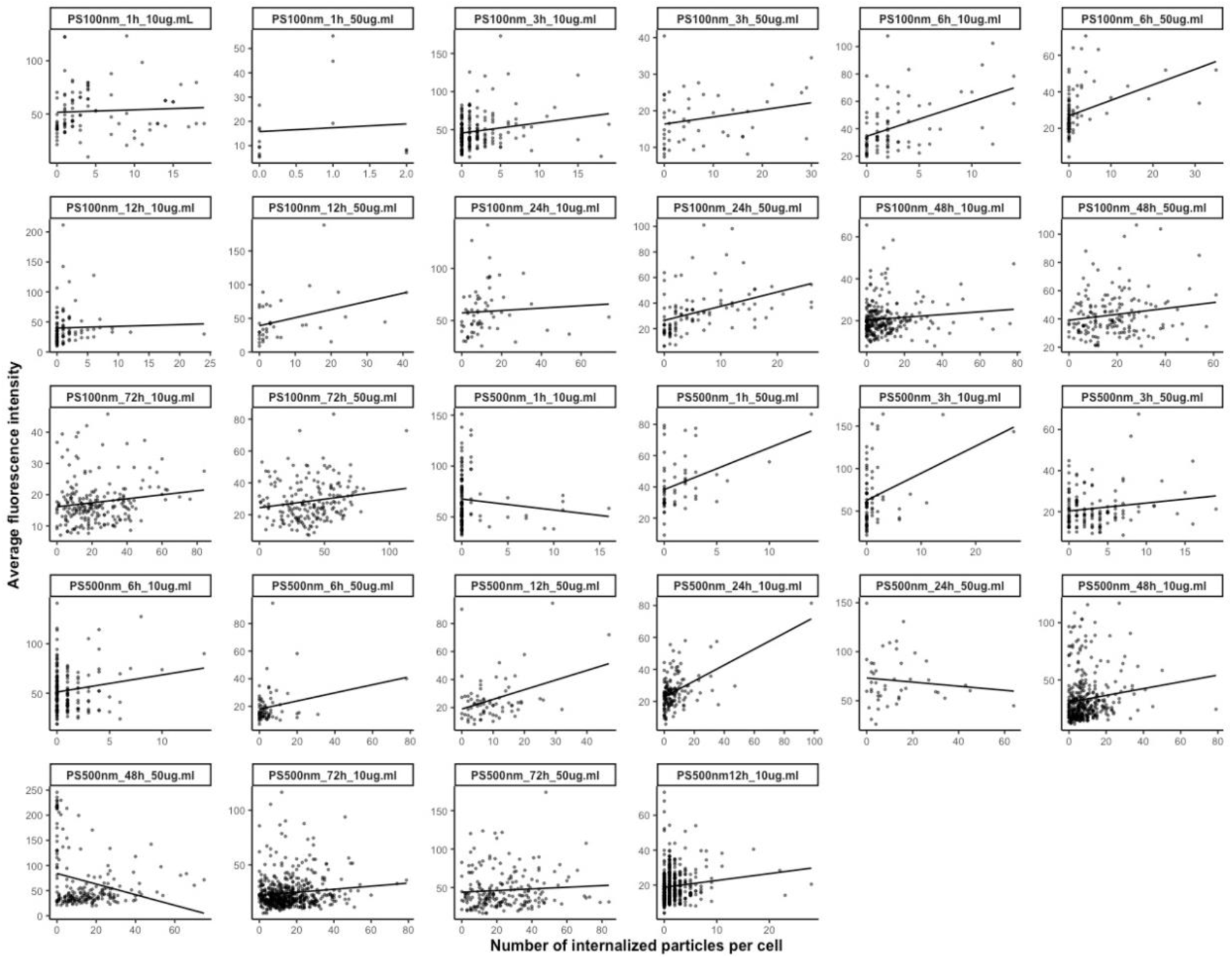
Scatter plots showing the correlation between the number of internalized PS-NPs per cell and the average fluorescence intensity of Iba1 in each cell after different exposure time points (1–72 h followed by washout steps) to PS-NPs of 100 nm or 500 nm at 10 or 50 µg/mL, as indicated. Represented data are derived from Pearson’s correlation coefficients (R) analysis.

### 3.8. IL-6 Expression

The obtained data showed transient IL-6 expression in BV2 cells, and various expression observed in different conditions. A slight increase in IL-6 levels was noted at 48 h, although the effects were not significant (Fig 12). The concentration, time and particle size also affected IL-6 expression.

**Figure 12.**
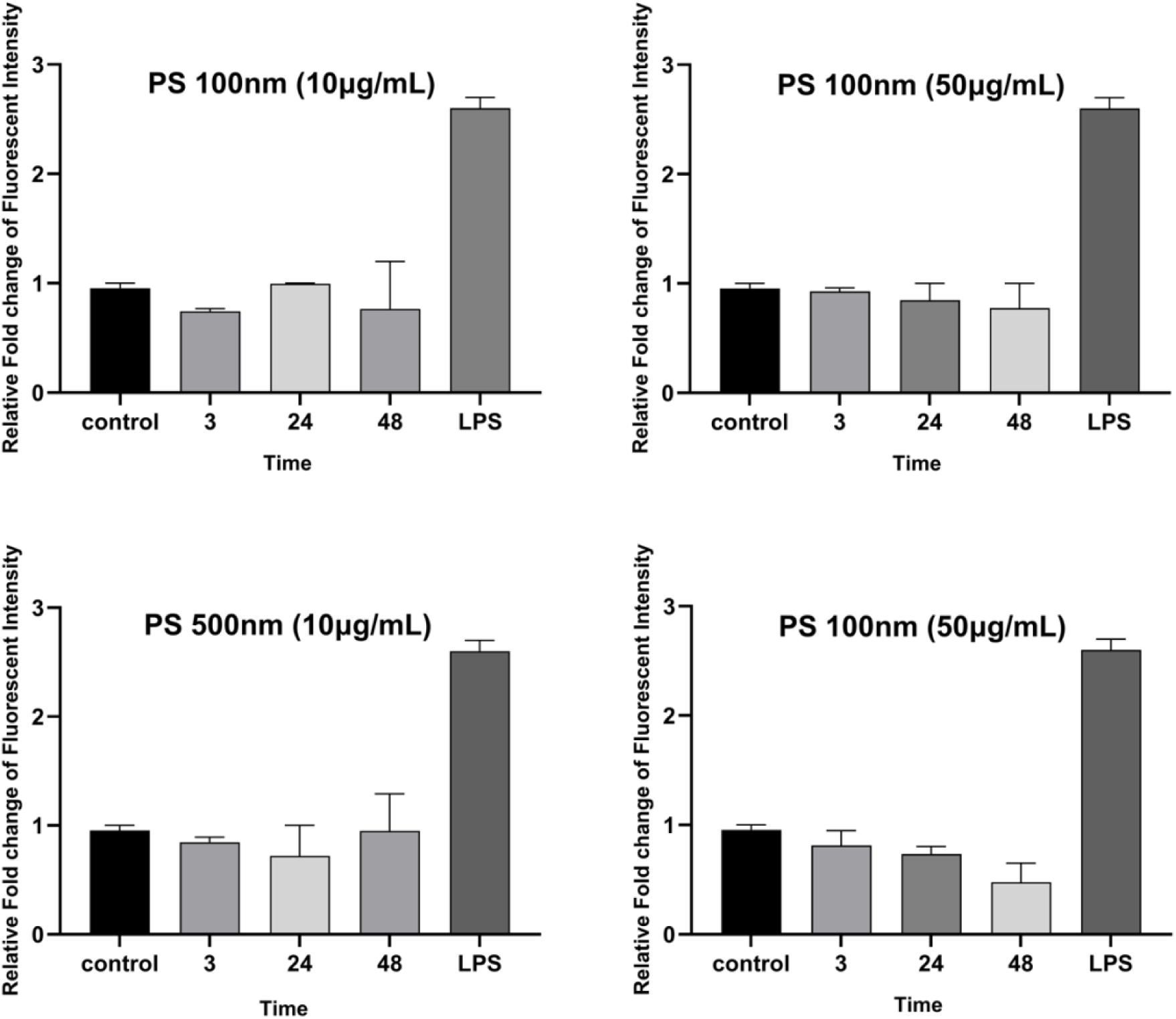
The IL-6 expression is depicted in this graph in BV2 cells treated with PS-NPs in different concentrations and times of exposure. The fluorescent intensity of IL-6 measured in different samples and controls. LPS was used as a positive control to detect IL-6 expression in BV2 cells. Data are represented as Mean ±SEM. One-way ANOVA with Dunnett’s post-test were used for statistical analysis. No significant changes were observed.

## 4. Discussion

There is a growing concern regarding the neurotoxic effects of nanoplastics [27]. In the complex composition of central nervous system, glial cells and more specifically, microglia, have important roles in brain homeostasis through their phagocytic activity and immune functions [28]. Shan et al. showed that 50 nm PS-NPs can cross the BBB and activate microglia in the brain of mice within 7 days of exposure. This was associated with an increased permeability of the BBB and neuronal damage [12]. Other reports have also suggested that NPs can cross the BBB or accumulate in the brain by other routes, such as olfactory translocation. Once in the brain, NPs may infiltrate various cell types including neurons, astrocytes, and microglia [29]. In our study, PS-NP exposure did not produce significant overt changes in the histological morphology of the brain, including the cerebral cortex. This is consistent with previous studies that showed environmentally relevant concentrations of PS-NPs often induce subtle molecular and cellular changes, particularly in microglia, without producing gross structural damage detectable by routine histology [30, 31].

Transcriptomic analysis revealed a clear size-dependent divergence in cortical responses to PS-NPs, characterized by subtle microglial priming following 100 nm PS-NPs exposure and robust neuroimmune and synaptic dysregulation following 500 nm PS-NPs exposure. Integration of IPA upstream-regulator predictions with functional categorization of differentially expressed genes provides mechanistic insight into how PS-NPs affect the transition from early adaptive signaling to overt neuroinflammatory dysfunction. Many of the differentially expressed genes and upstream regulators identified in our study, such as TGFB1, CX3CR1, APOE, CNTF, and CLEC10A/MGL2 are well-established components of microglial activation, immune regulation, and neuron–glia communication [32–34]. The directionality of the transcriptomic alterations aligns with our *in vivo* microglial morphology data and *in vitro* BV2 activation profiles, supporting a mechanistic link between PS-NPs exposure, microglial activation, and cortical gene expression changes.

Exposure to 100 nm PS-NPs showed a transcriptional state characterized by early microglial activation coupled with metabolic and xenobiotic adaptation rather than overt neurotoxicity, coordinated by the activation of AHR, HNF1A, INSR, and LHX1 upstream regulators. These regulators were strongly related to genes involved in vesicular trafficking and molecular transport such as *SLC18A2, SLC22A1, LCN2*, as well as immune and inflammatory signaling including *CCL5, IL22/IL22R2, ITGAD, CALCA*.

These observations support a role for microglia as primary targets to nanoplastic exposure, as previously reported [35]. Such transcriptional priming is consistent with the subtle microglial morphological remodeling observed *in vivo* and the transient activation profile measured in BV2 cells. Importantly, inhibition of CREB1 signaling in this condition suggests early disruption of trophic and homeostatic signaling pathways, which may reduce neuronal support and synaptic plasticity even in the absence of strong inflammatory gene induction [36, 37].

In contrast, the large increase in both the number and magnitude of differentially expressed genes in 500 nm PS-NPs exposed group, compared to 100 nm exposure group, indicate that larger particles result in a stronger biological response. The 500 nm PS-NPs exposure condition supports the conclusion that larger particles induce different neuroimmune and regulatory pathways. Previous in vitro studies using rat neuronal stem cells also observed a difference in DEGs between cells exposed to 50nm versus 500 nm PS-NPs [38].

CLEC10A/MGL2 (C-type lectin domain family 10 member A), also known as Mgl2 (macrophage galactose-type lectin 2), is a pattern-recognition receptor and a well-established marker of alternatively activated microglia, which appeared as a predicted activated node in the IPA network [39]. Its activation in our dataset suggests that, alongside pro-inflammatory signatures, cortical microglia may also engage compensatory or regulatory pathways involved in tissue remodeling, phagocytic clearance, and resolution-phase immune responses, confirmed by differentially expressed genes such as *ASGR1* and *GM5805*. Moreover, upstream regulator analysis identified inhibition of key neurotrophic and regulatory factors such as BDNF, AR, and TP73, all of which play critical roles in neuronal survival, cytoskeletal organization, and immune modulation [40, 41]. The suppression of these regulators was associated with altered expression of innate immune and inflammatory markers (*ZBP1, Ccl5, Cxcl10, Il8*) alongside remodeling-related transcripts (*MMP28, TAGLN, NOTCH1/3*) and downregulation of genes involved in neuronal maintenance and synaptic signaling (*SYN1, STX1B, SPTBN4, MAP1B, GRIN2B, ITPR1/2*) [42]. Also, the elevated expression of (*POMC*) as common differentially expressed gene in all conditions, suggests activation of stress-responsive neuroendocrine pathways following PS-NP exposure. As a precursor of ACTH and α-MSH, POMC plays a central role in hypothalamic–pituitary–adrenal (HPA) axis regulation and neuroimmune modulation, indicating potential involvement of integrated neuroendocrine–immune responses [43]. These observations are consistent with our findings of pronounced microglial branching simplification and *in vitro* evidence of size-dependent microglial activation. Direct comparison between 500 and 100 nm PS-NP exposure further highlighted selective suppression of microglial homeostatic regulators, including TGFB1 and CX3CR1. These molecules are central to microglial–neuron communication, synaptic pruning, and immune restraint [32]. Predicted inhibition of these pathways suggests that larger particles more effectively disrupt regulatory feedback mechanisms that normally limit neuroinflammatory signaling, as reflected by altered expression of microglial interaction and immune-associated genes such as *CCL5, ITGAD*, and *LCN2*, together with downregulation of pathways involved in synaptic signaling and neuronal membrane excitability [44, 45]. In contrast, predicted activation of CLEC10A/MGL2 (Mgl2), CNTF, and APOE indicates engagement of compensatory and regulatory microglial programs alongside inflammatory signaling [32–34]. These regulatory networks were associated with altered expression of genes involved in immune modulation, metabolic regulation, and cytoskeletal remodeling, including *LCN2, CCL5, ITGAD, PPARA, TAGLN, MYH9*, and *MYO5A*, consistent with a shift toward reactive and metabolically adaptive microglial states under conditions of increased particle burden [45]. Gene ontology analyses reinforced these findings by revealing widespread transcriptional alterations across biological processes involved in neuroinflammation, microglial activation, cellular junctions, cell signaling, neuronal plasticity, synaptic function, and cellular stress and metabolism. Since microglia regulate synaptic refinement, neuronal architecture, and extracellular signaling [46], coordinated suppression of these gene families likely reflects impaired microglial support functions and neuronal toxicity. Together with the absence of gross histopathological changes, these transcriptomic alterations suggest that PS-NP exposure induces a molecular vulnerability state in the cortex, characterized by weakened neurotrophic support and altered neuroimmune communication. Importantly, these transcriptomic findings correspond with the cellular mechanisms identified in this study. In our observations of *in vivo* exposure to PS-NPs, glial cells including microglia and astrocytes demonstrated activated phenotypes which reflect the inflammatory effects of PS-NPs exposure. Our data is consistent with previous studies including Paing et al. who showed that exposure to PS-NPs induced microglia and astrocyte activation [9]. Using a similar approach, Asmaa et al. also showed microglia and astrocyte activation in response to PS-NPs by assessing Iba1 and GFAP markers in mice models [47].

To further understand the mechanism of action of PS-NPs on microglia, we used BV2 cells which are a mice microglial cell line. Our data showed that BV2 cells showed transient but time dependent increase in Iba1, CD68, and IL-6 expression, thus support our *in vivo* observations in the cortex of mice chronically exposed to PS-NPs.

While the *in vivo* studies are essential parts of the toxicity study, having mechanistic assessments of nanoplastic internalization and activation of microglial cells are crucial contributions to our understanding of nanoplastic toxicity. Our *in* vitro data using BV2 cells exposed to PS-NPs and washout experiments provide a novel overview on nanoplastic toxicokinetics. Using the same cell type, several other studies have demonstrated microglial activation in response to nanoplastic exposure by assessing the expression of Iba1, and CD68 markers [7, 12]. Although the current scientific findings demonstrate microglial activation following PS-NP exposure, the realistic temporal dynamics of nanoplastic interactions with brain immune cells have been largely overlooked, as most studies rely on continuous exposure paradigms that do not reflect fluctuating or transient environmental exposures. To address this gap, we implemented an exposure–washout strategy to evaluate whether microglial responses persist after PS-NP removal. This approach is supported by emerging nanoparticle toxicology frameworks showing that cells may retain internalized particles for extended periods due to inefficient lysosomal clearance, leading to prolonged or delayed biological responses [48]. Sustained size dependent localization of PS-NPs in Rab7- and 100 nm PS-NPs in LAMP2-positive compartments suggest a mechanism by which these particles may contribute to long-term gene expression changes. Endolysosomal stress and impaired trafficking are known to influence microglial signaling, metabolic state, and cytokine responsiveness, thereby linking particle accumulation to long-term gene expression changes [49]. The convergence of morphological, functional, and transcriptomic data supports a model in which PS-NPs, particularly larger particles, drive size-dependent neuroimmune reprogramming through sustained intracellular retention rather than acute cytotoxicity. While the exposure is subchronic, the gene expression data may display greater changes as well as more prominent morphological effects following longer exposure periods. The transcriptomic data support the notion that nanoplastic neurotoxicity may be manifested primarily as chronic dysregulation of neuroimmune homeostasis, rather than immediate neuronal loss [50]. This has important implications for environmental risk assessment, given that such molecular and cellular alterations may predispose the brain to increased vulnerability under aging, secondary insults, or neurodegenerative conditions. On the other hand, PS-NPs uptake by BV2 cells in our *in vitro* study validates the capacity of PS-NPs to enter microglia. In addition, co-localization data suggest receptor-mediated endocytosis and lysosomal accumulation of PS-NPs. This observation corroborates results from Yu et al. (2025), who reported that nanoplastics accumulate in the brain of mice and are predominantly phagocytosed by microglia [13]. With respect to the type of plastic polymer, PS-NPs of 50-100 nm and MPs of up to 2 µm have been shown to enter the microglia, as demonstrated by *in vitro* studies in mouse-derived microglia cells [9, 14, 51]. Our observations of endolysosomal entry of PS-NPs, are also consistent with several studies, including Yin et al. [52], who showed that 100 nm PS-NPs entered mouse endothelial cells of the brain microvascular via the endolysosomal pathway [49]. A similar study by Liu et. al, in which rat basophilic leukemia (RBL-2H3) cells were exposed to 500 nm PS-NPs, demonstrated internalization of PS-NPs and significant accumulation in lysosomes [53]. In addition, Ding et, al. reported that RAB7 (late endosomes) and LAMP1 (lysosomes) levels were upregulated in PS-NPs-treated GES-1 cells (immortalized human gastric epithelial cell line), suggesting progression toward the late endosome and lysosome [54]. The same tendency is observed for other sizes of PS-NPs, including 50 nm that induced endolysosomal uptake in human neuronal cells [55]. Interestingly, within the same cell type, a similar pattern of internalization was observed when BV2 cells were exposed to PS-NPs of 80 nm [12]. However, the uptake kinetics or tendency for exocytosis of PS-NPs have not been well established. Similar to our present study, Liu et, al. reported that PS-NPs internalization increases rapidly in RBL-2H3 cells and was concentration and size-dependent. The phagocytic capacity of microglial cells is an important factor influencing their uptake ability [53]. Nanoplastics may enter microglia through phagocytic and endolysosomal mechanisms; however, since they cannot be degraded, they persist within the cells, consistent with our observations of long-term intracellular retention [51]. Together, these findings indicate that PS-NPs persist within microglial endolysosomal compartments, providing a mechanistic basis for sustained post-exposure effects including molecular and transcriptional signature. These alterations are essential in brain functions, since microglia and astrocytes play an important role in maintaining neuronal homeostasis [56]. Our findings revealed transient microglial activation even after washout, suggesting that PS-NPs exert a cumulative intracellular burden rather than an immediate, short-lived stimulus. Mechanistically, this is in line with the known behavior of nanoparticles that accumulate within late endosomes and lysosomes compartments characterized by slow turnover, limited degradation capacity, and susceptibility to trafficking dysfunction [54]. It is worth noting that the observed microglial activation occurred in the absence of sustained cytotoxicity. This observation would shed light on adaptive microglial scavenging responses to PS-NP exposure that may contribute to neuroinflammatory dysregulation. This distinction is critical, as microglial-mediated neuroinflammation can arise independently of cytotoxicity and may persist under sublethal stress conditions [9]. Consistent with this interpretation, the restrained cytokine response in our data suggests that PS-NPs trigger a controlled neuroimmune activation rather than a full pro-inflammatory cascade and such low-grade or transient cytokine signaling is characteristic of microglial priming states, which may sensitize the brain to secondary insults or prolonged stress without producing acute neuroinflammation [57, 58]. On the other hand, stronger correlations between particle burden and Iba1 expression at intermediate exposure durations suggest the existence of an activation threshold, beyond which microglia initiate cytoskeletal remodeling and functional responses. The subsequent decline in correlation at later time points, despite continued particle retention, is consistent with adaptive regulation or partial adaption mechanisms, consistent with the observed pattern of Iba1 expression in our study [59].

These findings provide a mechanistic bridge between PS-NP uptake kinetics, endolysosomal accumulation, transient microglial activation, and the transcriptional remodeling observed in cortical tissue, reinforcing the concept that nanoplastic-induced neuroimmune effects are governed by intracellular processing rather than particle presence alone.

## 5. Conclusions

This study provides integrated *in vivo* and *in vitro* evidence that environmentally relevant PS-NPs can reach the cortical region of murine brain and accumulate in microglial cells, where they cause transient activation and induce alterations in dynamics of the neuroimmune system without causing significant cytotoxicity or gross histopathological damage. Chronic oral exposure resulted in the cortical presence of PS-NPs and pronounced microglial morphological remodeling, highlighting microglia as a cellular target of nanoplastic neurotoxicity.

From a mechanistic point of view, PS-NPs were taken up by microglial cells via an endolysosomal pathway and exhibited prolonged retention in intracellular compartments, including late endosomes and lysosomes. This persistent intracellular burden of PS-NPs was associated with transient microglial activation, which was correlated with PS-NPs size, concentration, and duration of exposure, as reflected by Iba1, and CD68 expression. Interestingly, this study shows evidence for the first time that microglial activation is correlated with intracellular particle burden rather than cumulative exposure alone, supporting a threshold-driven and adaptive activation model, rather than continuous inflammatory escalation. Importantly, the observed data are in accordance with the transcriptomics analysis of the cortex, which showed size-dependent disruption of neuroimmune regulatory networks, including suppression of microglial homeostatic and neurotrophic pathways as well as activation of stress-adaptive signaling programs. These molecular alterations occurred without evidence of neuronal loss, suggesting that PS-NP exposure induced a state of cortical molecular vulnerability driven primarily by altered microglial function. Our findings emphasize that nanoplastic neurotoxicity occurred not solely by particle presence, but by intracellular processing, endolysosomal retention, and adaptive neuroimmune responses. Therefore, this work reinforces the importance of considering exposure dynamics, particle size, and post-exposure persistent effects in nanoplastic risk assessments and highlights microglia-concentrated mechanisms as critical determinants of long-term brain susceptibility to environmental nanoplastics.

## 6. Acknowledgements

The authors thank Dr. N Sonenberg and R. Ladak for assistance with the BV2 cells (McGill University). J. Tremblay is thanked for his assistance with both confocal microscopy and flow cytometry. This study was supported by an NSERC Alliance Grant (ALLRP 558452 – 20) (SL and DGC) and NSERC Discovery grant (RGPIN-2021-03330) (SC) and a Canada Research Chair in Reproductive Toxicology to DGC.

## 7. CRediT authorship contribution statement

All of the authors have contributed to the planning of different experiments, sampling, analysis, interpretation and writing of the manuscript.

## 8. Declaration of competing interest

The authors declare that there is no financial or non-financial conflict of interest.

## Notes

### Competing Interest Statement

The authors have declared no competing interest.

