## Supplemental figure for "Polystyrene Nanoplastics Accumulate in Murine Cortex and Induce Transient Microglial Activation via Endolysosomal Retention"

Supplemental Figure 1

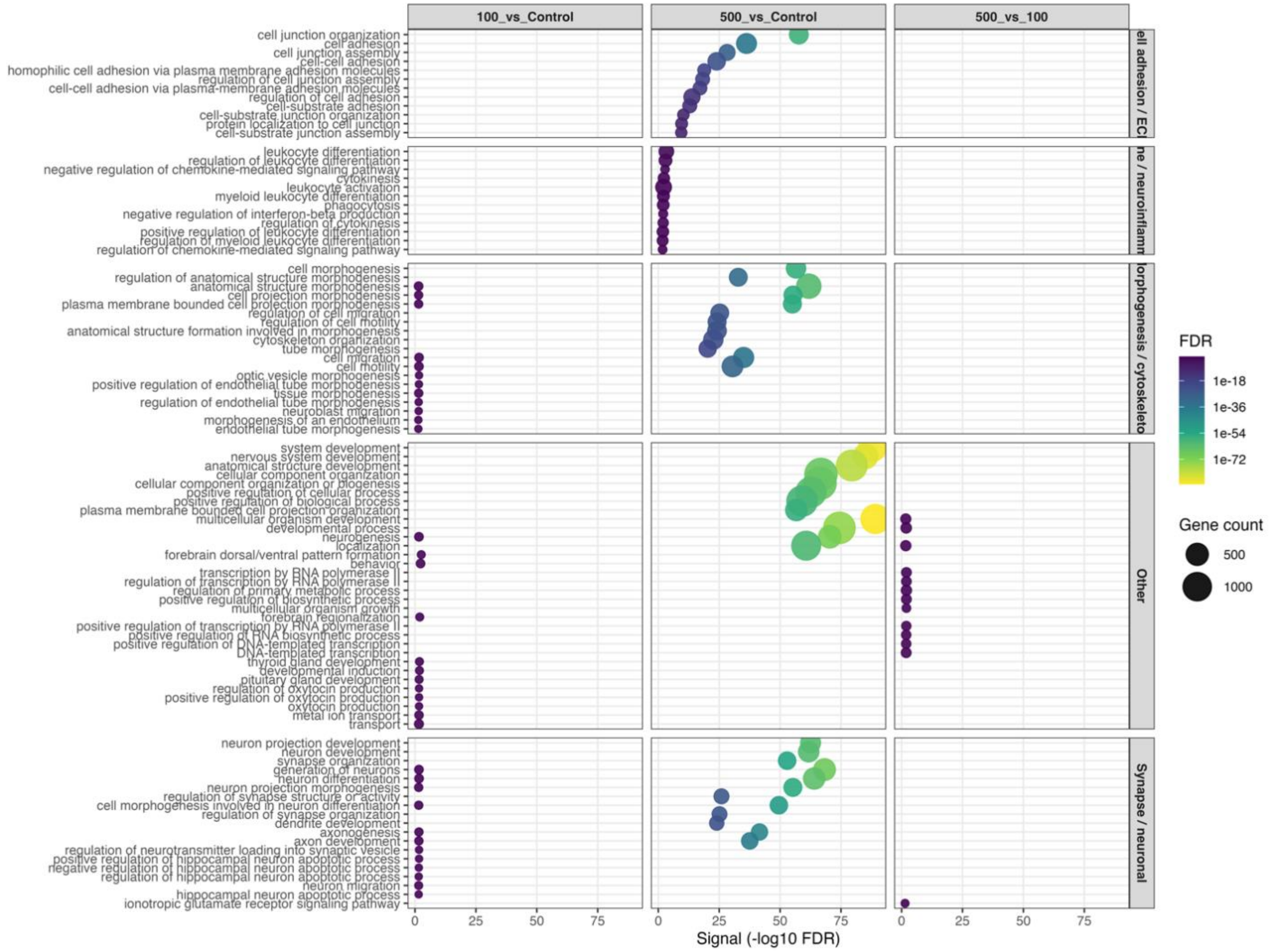


Supplementary figure 1. Comparative Gene Ontology (GO) enrichment analysis across exposure conditions. Bubble plots demonstrate significantly enriched GO Biological Process terms identified in cortical tissue following PS-NP exposure. Panels represent the three pairwise comparisons (100 nm vs Control, 500 nm vs Control, and 500 nm vs 100 nm). Bubble size corresponds to the number of genes contributing to each GO term (gene count), and bubble color indicates enrichment significance (FDR-adjusted p-value). The analysis reveals a pronounced enrichment of immune-related, membrane-associated, synaptic, and neuronal development processes, with stronger and broader enrichment observed in the 500 nm vs Control comparison, highlighting size-dependent transcriptional effects of nanoplastic exposure.
